# Differential phase register of Hes1 oscillations with mitoses underlies cell-cycle heterogeneity in ER+ breast cancer cells

**DOI:** 10.1101/2021.01.04.425227

**Authors:** Nitin Sabherwal, Andrew Rowntree, Elli Marinopoulou, Tom Pettini, Sean Hourihane, Riba Thomas, Jochen Kursawe, Ximena Soto, Nancy Papalopulu

## Abstract

Here, we study the dynamical expression of endogenously labelled Hes1, a transcriptional repressor implicated in controlling cell proliferation, to understand how cell-cycle length heterogeneity is generated in ER+ breast cancer cells. We find that Hes1 shows oscillatory expression with approximately 25h periodicity and during each cell-cycle has a variable peak in G1, a trough around G1-S transition and a less variable second peak in G2/M. Compared to other subpopulations, the cell-cycle in CD44^High^CD24^Low^ cancer stem cells is longest and most variable. Most cells divide around the peak of the Hes1 expression wave but preceding mitoses in slow dividing CD44^High^CD24^Low^ cells appear phase-shifted, resulting in a late-onset Hes1 peak in G1. The position, duration and shape of this peak, rather than the Hes1 expression levels, are good predictors of cell-cycle length. Diminishing Hes1 oscillations by enforcing sustained expression slows down the cell-cycle, impairs proliferation, abolishes the dynamic expression of p21, and increases the percentage of CD44^High^CD24^Low^ cells. Reciprocally, blocking the cell-cycle causes an elongation of Hes1 periodicity, suggesting a bidirectional interaction of the Hes1 oscillator and the cell-cycle. We propose that Hes1 oscillations are functionally important for the efficient progression of the cell-cycle and that the position of mitosis in relation to the Hes1 wave underlies cell-cycle length heterogeneity in cancer cell subpopulations.

**Significance statement:** Tumours exhibit heterogeneities that are not due to mutations, including Cancer Stem Cells with different potencies. We show that the cancer stem cell state predisposed to dormancy *in vivo* has a highly variable and long cell-cycle. Using single-cell live-imaging for the transcriptional repressor Hes1 (a key molecule in cancer), we show a new type of circadian-like oscillatory expression of Hes1 in all cells in the population. The most potent cancer stem cells tend to divide around the trough of the Hes1 oscillatory wave, a feature predictive of a long cell-cycle. A novel concept proposed here is that the position of cell division with respect to the Hes1 wave is predictive of its prospective cell-cycle length and cancer cellular sub-state.

## Introduction

Molecular and phenotypic analyses at the single-cell level have revealed a great deal of heterogeneity in genetically identical populations of cells, which were presumed to be homogeneous based on the population averaging methods. Such non-genetic heterogeneity is thought to have several potential sources, ranging from transcriptional noise (intrinsic) to variability of exposure to environmental signals (extrinsic). Non-genetic heterogeneity has both benefits and pitfalls for the optimal function of biological systems (1). In cancer, it represents a significant challenge because non-genetic heterogeneity is a crucial factor underlying the differential response to chemotherapy, the emergence of treatment resistance in the absence of new mutations, as well as the disease relapse driven by the reactivation of dormant cancer stem cells (2) (3).

At the phenotypic level, non-genetic heterogeneity can manifest itself in the form of the cell-cycle length heterogeneity, which is particularly important in cancer as it may underlie the transition between rapidly dividing, slowly dividing and quiescent cancer cells, with implications for cancer progression, relapse and development of resistance to the treatment (4). Thus, there is a need to understand how cell-cycle heterogeneity may be generated within a population of cells at a mechanistic level.

Most molecular studies of heterogeneity have relied on snapshot measurements of populations of cells where it can be difficult to distinguish the contribution of dynamical gene expression in the time domain versus different levels of expression which could be relatively stable over time. The contribution of dynamical gene expression in creating heterogeneity has been brought to the forefront by studies showing that important regulatory molecules show oscillatory expression (5) (6). Oscillations in gene expression that are asynchronous between cells are not apparent when viewed as a snapshot and on the whole, are an underappreciated source of non-genetic heterogeneity in a population of cells. The ever-growing list of such molecules includes genes associated with the cell-cycle and the circadian clock but also genes such as the DNA damage response protein p53, the pro-neural protein Ascl1 and the neurogenic ligand Dl, to mention just a few (6). Notable examples are the key transcription factors (TF) of the Hes/Her (mammals/zebrafish) family of genes, which have been shown to oscillate synchronously in somitogenesis and asynchronously in neural progenitor cells and whose oscillations enable the cell-state transition to differentiation (7) (8) (9) (10) (11). Among Hes family genes, Hes1 is of particular importance because it has been implicated in controlling proliferation in a cancer context and shown to control (directly or indirectly) the expression of genes involved in cell-cycle. Indeed, Hes1 is known to affect cell-cycle by repressing both activators (CyD, CyE and E2F) and inhibitors (p21 and p27) of cell-cycle, directly or indirectly (12) (13) (14) (15) (16). However, how these conflicting activities of Hes1 are incorporated in the cell-cycle machinery to modulate its kinetics are not well understood. It is also not clear whether Hes1 oscillates in a cancer model and if so, how does the oscillatory expression of Hes1 interface and possibly control the cell-cycle.

Here, we use an endogenously tagged reporter to ask whether Hes1 oscillates in ER+ breast cancer cells and if so, whether it shows a reproducible correlation with the cell-cycle. We have chosen to focus on MCF7 cells, a key cellular model system of ER+ breast cancer because it is a type of cancer with long term relapse, a known clinical problem (17), which may occur at least in part due to non-genetic heterogeneity, cell plasticity and exit from quiescence at the cellular level. MCF-7 cells are a good model to look at the influence of gene expression dynamics in ER+ breast cancer because they contain well characterised subpopulations of cancer stem cells (ALDH^High^ and CD44^High^CD24^Low^ cells), which differ in their ability for tumour re-initiation and over time, convert into each other (18).

We report that Hes1 oscillates with an average of 25hr periodicity and this nearly matches the average cell-cycle length of ~24hr in most of the MCF7 cells. Through clustering analysis of single-cell dynamic traces of Hes1 over the cell-cycle, we find a reproducible relationship between the Hes1 dynamics and the cell-cycle, such that in most cells, the preceding division takes place at or near the peak of Hes1 expression. This peak in Hes1 protein expression is then followed by a dip, the onset of which is followed by the G1-S transition in most cells, leading to a second period of increased Hes1 protein concentration before the next division. A minority of the cells show a delayed and prominent Hes1 peak further into the G1 phase, making them appear as if they divided at the trough rather than the peak of Hes1 expression. In such cells, the division can be described as “out of phase” with the Hes1 oscillator. Such cells tend to have longer cell-cycles, and remarkably, they are enriched for CD44^High^CD24^Low^ cancer stem cells, which also show the highest cell-cycle heterogeneity and the highest propensity to switch in or out of phase. Finally, when we experimentally dampened the Hes1 oscillations, the cell-cycle slowed down, p21 dynamics were abolished and the proportion of CD44^High^CD24^Low^ increased, indicating the functional significance of Hes1 oscillations for an efficient cell-cycle progression and cell fate. Conversely, blocking the cell-cycle with CoCl_2_ elongated the periodicity of Hes1, suggesting a bidirectional coupling of the Hes1 oscillator and the cell-cycle.

We conclude that Hes1 dynamics are important for sculpting the cell-cycle, such that it occurs efficiently and with little heterogeneity, around a set average cell-cycle length. We propose that differential alignment of Hes1 dynamics with mitoses (i.e. the phase registration) underlies cell-cycle heterogeneity in breast cancer cells and may underlie the propensity of some stem-like cancer cells to become quiescent.

## Results

### 1. CRISPR/Cas9 mediated N-terminal tagging of endogenous Hes1 with mVenus in ER+ breast cancer cells (MCF7)

To study Hes1 expression dynamics in ER+ breast cancer MCF7 cells at single-cell level and in real-time, we generated an in-frame fusion of mVenus with the endogenous Hes1. A cDNA cassette expressing mVenus was N-terminally fused to the first exon of Hes1 using a CRISPR/Cas9 genome editing approach (homology-directed repair/HDR), followed by FACS-based clonal selection and expansion of mVenus+ single cells (see materials and methods, m&m, Fig1A). After genotyping, a clonal line (hereafter named as Clone 19/Cl19) was chosen for any further analysis (FigS1 and movieS1). Cl19 cells were found to be hemizygous for Hes1, with one Hes1 allele having correct insertion of the mVenus cassette and another one knocked-out due to insertion of frame-shift mutation after CRISPR reaction. Cl19 line being hemizygous for Hes1 eliminated any possible interference to the tagged allele by the untagged allele. Cl19 cells recapitulated the endogenous nuclear Hes1 expression and spatial heterogeneity, as assessed by Hes1 immunostaining of parental MCF7 cells and mVenus-Hes1 snapshots from live-imaging of Cl19 cells (Fig1B-D). During Incucyte based image analysis, both parental MCF7 and Cl19 cells exhibited similar growth rates (FigS2), suggesting that knocking-out a Hes1 allele in Cl19 cells did not adversely affect the cells.

**Fig1.**
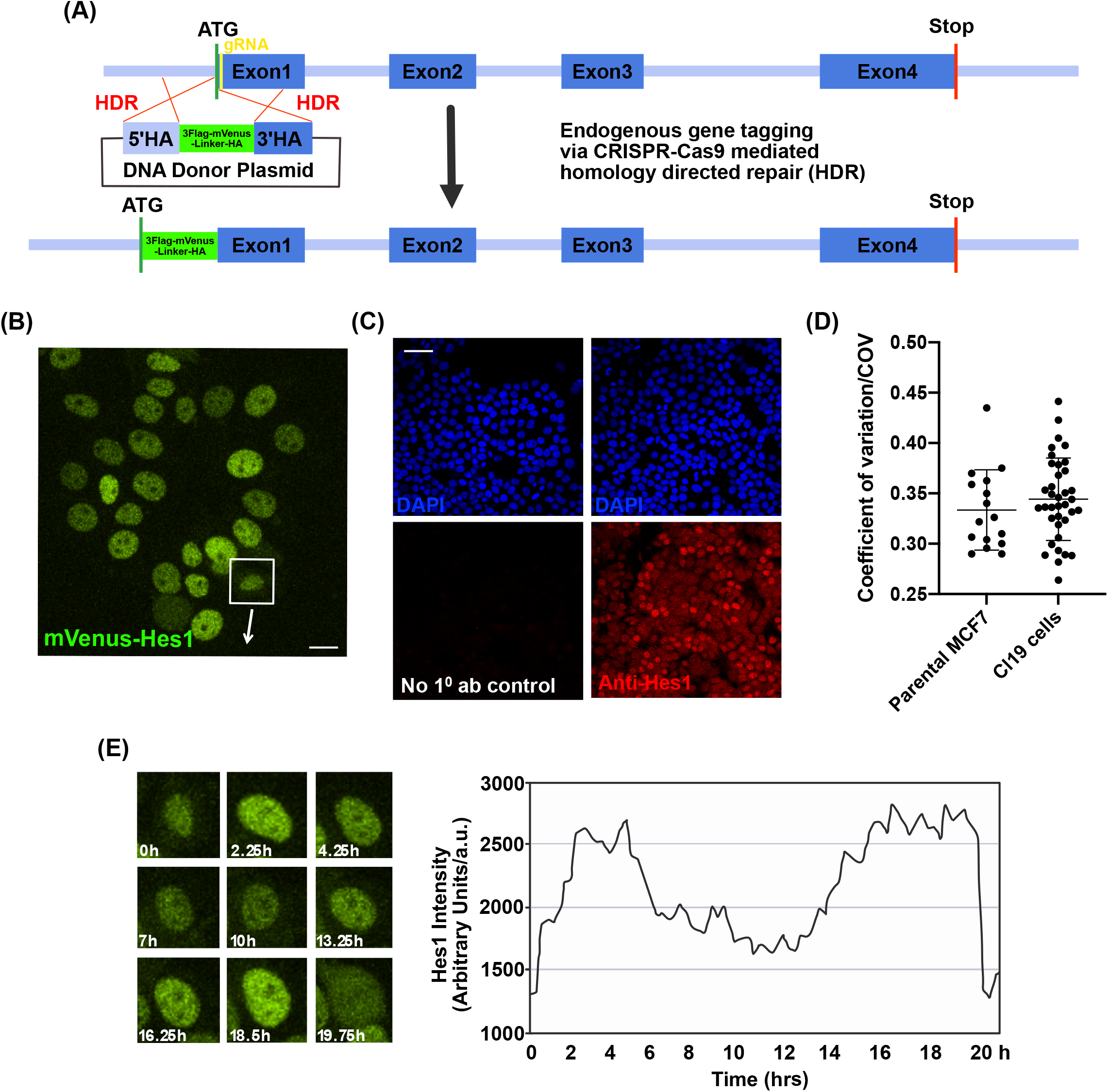
CRISPR-Cas9 mediated N-terminal tagging of the endogenous human Hes1 with mVenus in MCF7 cells shows snapshot heterogeneity due to temporal dynamics. (A) Schematic of the Hes1 genomic locus, before and after tagging, the donor plasmid, guide RNA and how CRISPR-Cas9 mediated homology-directed repair (HDR) reaction achieves in-frame tagging of the first exon of Hes1 with mVenus cDNA cassette. (B) Imaging of the sequence-verified clonal CRISPR-Cas9 line (Cl19) shows nuclear localisation of Hes1 (Scale bar=20um) and (C), snapshot heterogeneity in the level of endogenous mVenus-Hes1 expression, which were recapitulated by the immuno-staining of parental MCF7 cells using an anti-Hes1 antibody (Scale bar=50um). (D) Snapshot heterogeneity as measured by estimating coefficient of variation (COV) from images of Cl19 cells and Hes1-immunostained MCF7 cells. COV values (a measure of spatial heterogeneity) showed no differences between parental MCF7 cells and the CRISPR line (Cl19). (E) Example of a single cell (highlighted with a box in B) tracked over time. Time-lapse images of this example cell show that Hes1 protein levels fluctuate over time (left panel), resulting in a dynamic Hes1 expression as quantified by IMARIS-based cell-tracking (right panel).

We then compared protein half-lives of mVenus-Hes1 in Cl19 cells by live-imaging against an HA-tagged Hes1 transfected in parental MCF7 cells by western blot analysis, after Cycloheximide (CHX) treatment. Both the endogenous mVenus-Hes1 and the exogenous HA-Hes1 exhibited half-lives of around 4h, which is longer than the mouse protein (24-26min) (19), consistent with the overall increased protein stability in human cells (20). These results suggest that mVenus fusion to endogenous Hes1 does not affect its half-life (FigS3).

The snapshot heterogeneity of Hes1 expression (Fig1B and D) may originate (alone or in combination) from two possible scenarios that are, the presence of cells with various stable Hes1 expression levels and/or a temporally dynamic Hes1 expression, such as random fluctuations or oscillatory protein expression. Single-cell live-imaging over time suggested that the expression of Hes1 is temporally dynamic (Fig1E), which was analysed further in detail during this study (see below). To ensure the reproducibility of the dynamics in this clone, we compared it with two other CRISPR/Cas9 clones - one hemizygous (M18) and another heterozygous line (Cl12) and found them to have similar Hes1 dynamics (FigS4A-C). No difference in the snapshot heterogeneity in Hes1 expression between the reporter line (Cl19) and the parental MCF-7 cells also suggested that the reporter accurately reflects the endogenous dynamics (Fig1B-D). Thus, population heterogeneity of Hes1 protein expression in MCF-7 is due, at least in part, to temporal Hes1 dynamics, which may be oscillatory, as we assess in section 4 below. Furthermore, smFISH against Hes1 performed on parental MCF7 and Cl19 cells showed that number of mature Hes1 mRNA molecules are not constant through the cell-cycle, are higher in the G2 phase than in the G1 phase and suggested some dynamic Hes1 behaviour even at mRNA levels both in parental and CRISPRed MCF7 cells (FigS4D).

### 2. MCF7 cells exhibit high cell-cycle length heterogeneity, which is higher for the stem cells

MCF7 cells in culture are mainly representative of non-stem cells, constituting the majority of the population (~99%), referred to as ‘bulk’ cells. A tiny proportion of cells are classed as breast cancer stem cells (bCSCs) in terms of their functional roles in disease progression; CD44^High^CD24^Low^ bCSCs tend to be quiescent *in vivo*, while ALDH^High^ type bCSCs are proliferative (18). Therefore, to test whether stem cells might display interesting and useful deviations from the majority of the cells present in culture, we FACS-sorted Cl19 cells into three subpopulations (see m&m) for imaging and analyses. A similar number of cells from each subpopulation (ALDH^High^, CD44^High^CD24^Low^ and bulk cells) were compared against one another, or, if required, pooled together to generate a virtual ‘mixed’ population. Both unsorted and FACS-sorted cellular subpopulations from MCF7 cells are highly amenable to single-cell live-imaging experiments. We live-imaged subpopulations of cells from Cl19 line for durations encompassing at least one complete cell-cycle (mitosis to mitosis). To avoid any effects of synchronising agents on either the cell-cycle length or the Hes1 expression, we imaged non-synchronised cells at different stages of their cell-cycle. However, later, for data analyses and comparisons, we pseudo-synchronised Hes1 time-tracks, using the mitoses points as our landmarks.

When characterising the Hes1 levels and dynamics in relation to the cell-cycle, it is crucial to consider the subpopulation heterogeneity that exists in MCF-7 cells. We estimated the cell-cycle length from single-cell live-imaging data collected for subpopulations of Cl19 MCF7 cells. The reconstituted ‘mixed’ population data showed a mean cell-cycle length of 29.3h (SD = 11.8h, median = 25.8h, IQR = 9.6h) with high heterogeneity in the cell-cycle lengths with a heavy-tailed distribution towards longer cell-cycles (Fig2A). ‘Bulk’ cells which represent the majority of cells in the unsorted, heterogeneous cell population had a mean cell-cycle length of 23.4+/−6.6h (mean with SD). By contrast, both stem cell populations had significantly longer cell-cycle lengths, which were on average 29.3+/−10.5h for ALDH^High^ cells and 33.1+/−14.3h for CD44^High^CD24^Low^ stem cells (Fig2B). The numbers agree with the pre-existing notion that CD44^High^CD24^Low^ cells represent the quiescent-like stem cells (18), dividing the slowest in normal culture conditions, and ALDH^High^ type stem-cells are more proliferative. In addition, both stem cell populations exhibited higher levels of cell-cycle heterogeneity than the bulk cells, and this was the highest for the CD44^High^CD24^Low^ population, evidenced by the higher ‘spread’ of cell-cycle length values (CD44^High^CD24^Low^ interquartile rage/IQR = 15.2h, ALDH^High^ IQR = 8.25h and Bulk IQR= 5.3h) (Fig2B).

**Fig2.**
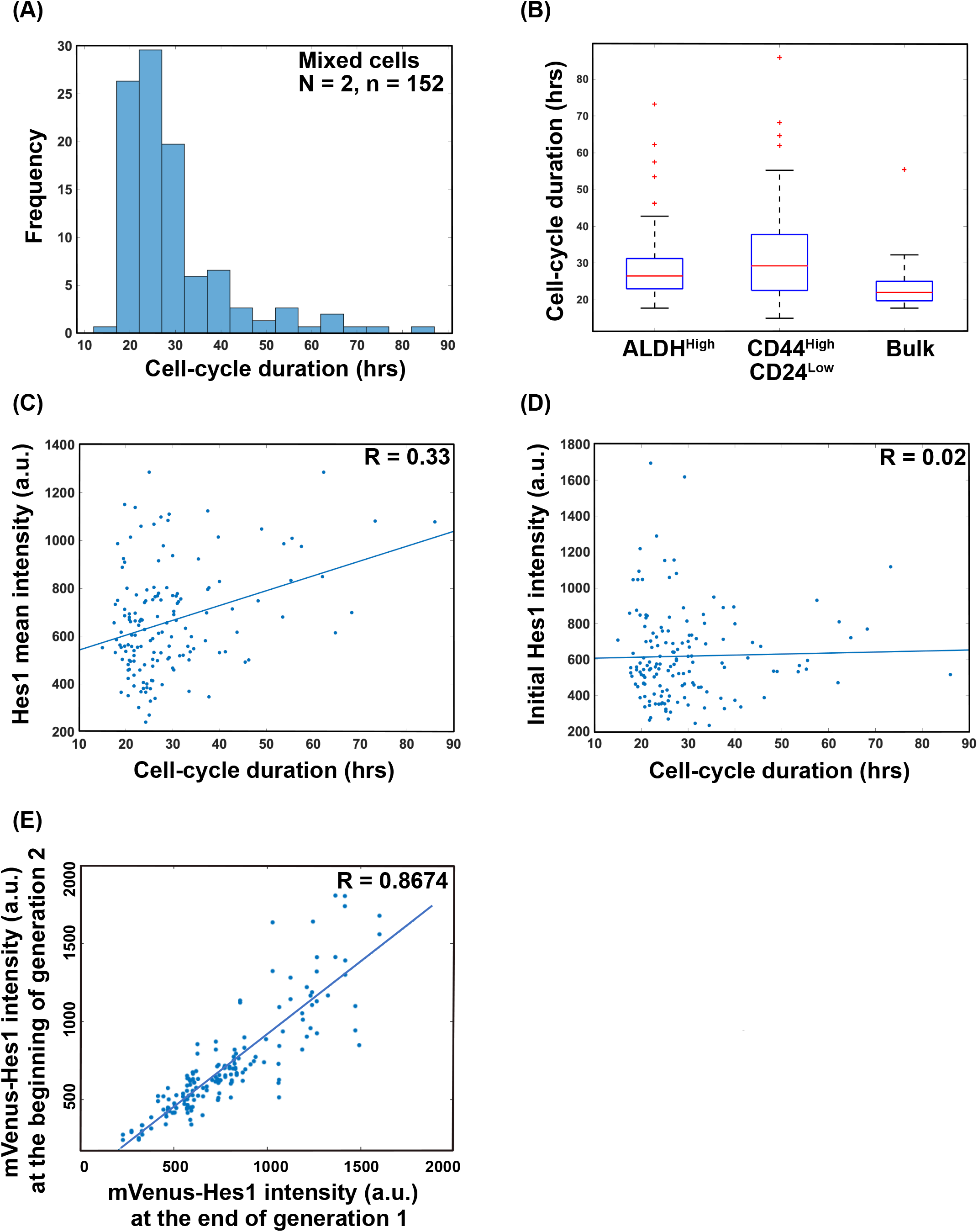
Cell-cycle length heterogeneity in MCF7 cells shows no relationship with the Hes1 mean levels. (A) Cell-cycle length of ‘mixed’ MCF7 cells has a mean value of around 29h but shows high heterogeneity (notice the heavy-tailed distribution of the cell-cycle length). (B) Cell-cycle is longer and more heterogeneous for stem cell subpopulations than bulk cells; CD44^High^CD24^Low^ cells showed the longest and the most heterogeneous cell-cycle. Heterogeneity in these cells was even higher than ALDH^High^ stem cells with Bartlett’s test showing that CD44^High^CD24^Low^ and ALDH^High^ cells have significantly different variances (*, p = 0.0192). (C) The correlation between the Hes1 mean intensity and the cell-cycle length was overall very modest (R = 0.33) as only 11% (R^2^ = 0.11) of cell-cycle length variance could be attributed to its correlation with the Hes1 levels. (D) The cell-cycle length also did not correlate with the initial Hes1 mean intensity in single cells (immediately after mitoses, in the earlier G1 phases). (E) Mean Hes1 levels do not change immediately prior to and after mitoses. Overall, these data suggest that Hes1 mean expression levels are not a dominant influence on the cell-cycle length and heterogeneity.

### 3. The cell-cycle length shows very weak correlation with the mean Hes1 level

Given the significant heterogeneities in cell-cycle lengths among MCF7 cells, we also asked if there is a correlation between Hes1 expression and the cell-cycle, that would imply a functional role. We first examined the correlation of Hes1 protein level with the cell-cycle length by plotting the mean Hes1 levels against cell-cycle lengths. A Spearman correlation coefficient (R) value of only 0.33 suggested only a very weak correlation between the two (Fig2C); the corresponding R^2^ value of 0.11 would mean that only 11% of the variance in the cell-cycle length can be attributed to its correlation with the Hes1 mean levels. Furthermore, we found no correlations between the cell-cycle lengths and the initial Hes1 mean intensity (immediately after mitosis) (R = 0.02) (Fig2D). We also noticed that the mean Hes1 expression in the beginning of the cell-cycle is pretty identical to the mean Hes1 expression at the end of the previous cell-cycle, meaning there is no sudden loss or gain of Hes1 protein over mitosis (Fig2E). This also suggested that the mean Hes1 levels towards the end of previous cell-cycle would not affect the next cell-cycle. We hence proceeded by analysing to what extent dynamics of Hes1 expression, rather than its overall expression level, have an influence on the cell-cycle.

### 4. Endogenous mVenus-Hes1 protein expression in MCF7 cells is oscillatory with a circadian-like periodicity

We investigated whether Hes1 shows oscillatory expression in MCF7 subpopulations as has been reported in many developmental systems (8) (9) (21) (22) (23) (24) (25). We ran mVenus-Hes1 time traces obtained by stitching together Hes1 dynamics over two consecutive cell divisions, through pyBOAT/Wavelet, a periodicity analyses platform (Fig3) (26). The power threshold values for the analyses of actual Hes1 traces (Fig3A-F) were set based on control background traces, which were analysed similarly but showed no periodicity (Fig3G-I). Fourier analysis showed that unlike the control traces (Fig3H), mVenus-Hes1 expression showed dominant periods (Fig3B and E), with mean period around 25h (Fig3J). In addition to the pyBOAT platform, we also ran our mVenus-Hes1 single-cell time trace data through Lomb-Scargle periodogram (LSP) (FigS5A-B) (27). As a biological negative control, we used a viral reporter line – Ubc:NuVenus (expressing mVenus tagged with SV40 nuclear localisation signal under the Ubc promoter) – which is not expected to exhibit oscillatory dynamics. LSP analysis showed a sharp peak on the power spectrum for mVenus-Hes1 traces but not for the control (Nu-mVenus) traces (FigS5B); the normalised power values were significantly higher for mVenus-Hes1 than for Nu-mVenus traces (FigS5C). This confirmed that the dynamics of Hes1 protein expression conferred the periodicity observed with the endogenous mVenus-Hes1 reporter. mVenus-Hes1 periodicity analyses with pyBOAT was further corroborated using Autocorrelation methods (28), showing a mean value of ~26h and similar distributions for Hes1 periodicity values (Fig3J).

**Fig3.**
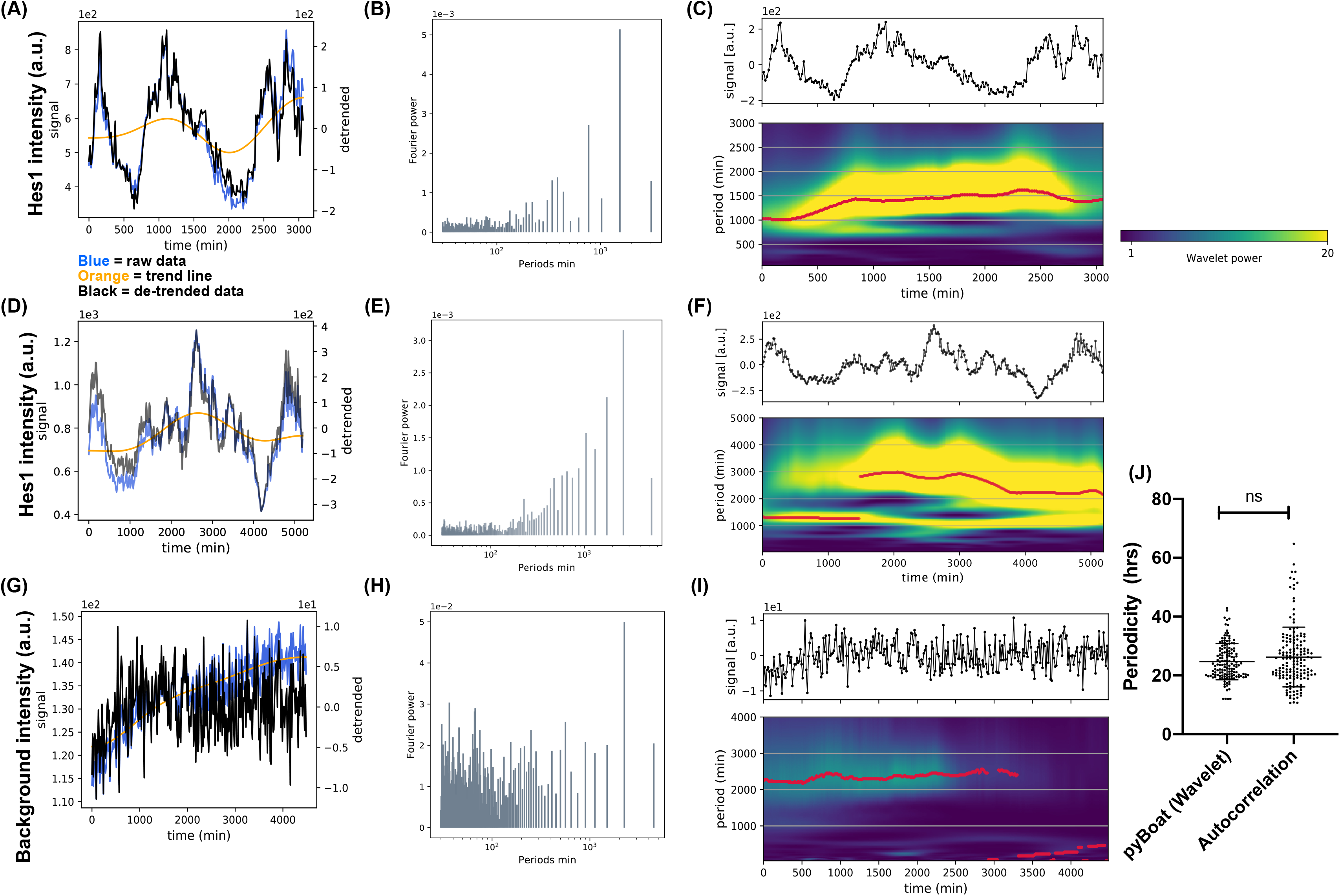
Wavelet analyses of mVenus-Hes1 single-cell time tracks from cellular sub-population showed evidence for Hes1 wave with a nearly 24hr periodicity. (A) Using pyBoat platform, mVenus Hes1 time series are detrended prior to periodicity analysis. The black lines show an example of a raw mVenus recorded time series. Blue is the corresponding detrended time series. The orange line denotes the trend. (B) Fourier power spectrum for the same input Hes1 trace shows a dominant period peak at around 25h. (C) The detrended mVenus-Hes1 trace is shown again in the above panel, with the lower panel showing the associated period-time-power heat map collapsed in 2D. The periods with the highest power at each time point are used to estimate the mean period and power across the duration of the input trace. (D-F) shows similar analyses of another Hes1 trace, which shows elongation of periodicity over time (see red ridge lines in F, as compared to red line in C). (G-I), Fourier and wavelet analysis of a background time series, with no dominant period for the entire track as confirmed through the Fourier power spectrum (F) and the power values at the strongest wavelet periods which are lower than in (C). Notice more than 10-fold higher Fourier power for Hes1 trace in comparison to the background trace (F *vs* B and E), and also highly coherent period-power values (bright yellow colour in C and F, shown as a ridge in red) for Hes1, compared to low, non-coherent period-power values for background trace (very low yellow content in G). (J) A mean period of around 25h was confirmed in the population through different analyses methods (Autocorrelation *vs* pyBOAT).

As the Hes1 periodicity we report here was higher than the previously reported Hes1 periodicity in mouse cells (ultradian oscillations of 2-3h) (23), we looked for evidence of ultradian periodicity in the Hes1 traces in MCF-7 cells using a Gaussian Processes-based method (29). Interestingly, an ultradian periodicity of 5-6h was observed which was the same in different cellular subtypes but this was embedded within the longer periodicity, giving rise to ‘nested oscillations’ (FigS6A-B). None of the features of ultradian oscillations (periodicity, fold change) correlated well with the cell-cycle length (FigS6C-D); therefore, we did not follow these further in context of the cell-cycle heterogeneity.

### 5. Dynamic Hes1 expression between two consecutive mitoses points is biphasic

Next, we asked whether the Hes1 dynamic expression has a reproducible relationship with the cell-cycle. To facilitate this and for the comparison of large numbers of dynamic Hes1 traces, we visualised all Hes1 traces simultaneously using heatmaps of expression from mitosis to mitosis (see m&m and FigS7). To highlight differences in Hes1 expression dynamics rather than their levels we normalised Hes1 traces using Z-scores [(raw intensity - mean intensity)/standard deviation], which enforced that each visualised trace has the same mean and variance. We organised these normalised heat maps in the increasing order of the cell-cycle lengths using second mitoses as a common end-point (Fig4A). We also aligned individual, normalised mVenus-Hes1 time series traces using second mitoses as a common end-point (Fig4D). Both these alignments showed a biphasic Hes1 expression among all cells, wherein a first Hes1 protein peak was followed by a dip and then by an ascending Hes1 expression towards the end of the cell-cycle (Fig4A and D). To ensure that this dynamic behaviour was specific to Hes1, we expressed Venus and nuclear Venus under the control of the Ubc constitutive promoter as controls (Fig4B-C). We analysed their expression traces in the same way as for the mVenus-Hes1. No deducible or reproducible dynamics were observed with either of these controls.

**Fig4.**
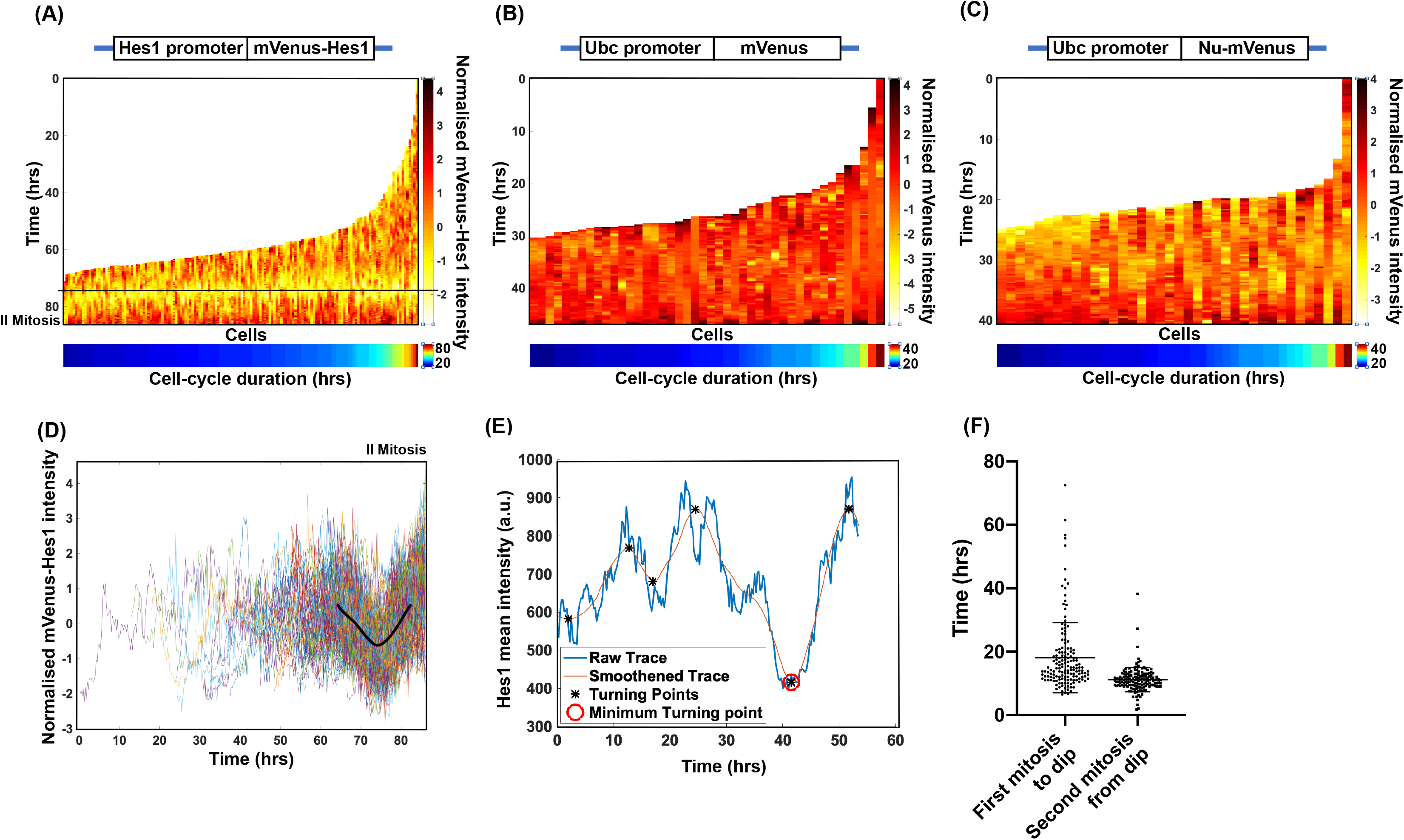
Dynamic Hes1 expression between two consecutive mitosis points is biphasic. (A) Endogenous mVenus-Hes1 traces from mixed Cl19 cells plotted as heat maps (above; see FigS7) and organised in increasing order of cell-cycle length (below). Notice the presence of the Hes1 expression dip (highlighted by a black line), preceded by a highly variable and followed by a less variable Hes1 peak, giving rise to the biphasic Hes1 profile during the cell-cycle. When mVenus (B) or Nu-mVenus (C) were expressed under the Ubc promoter as controls, their expression profiles during the cell-cycles showed no such characteristics. (D) When we plotted the same individual Hes1 tracks (as in A) as line plots, collapsed at the second mitoses at the right-hand side of the graph, we noticed a similar biphasic Hes1 expression dynamics with a variable, first Hes1 peak, followed by a dip in its expression (again marked by a black line), which was followed by a Hes1 peak towards the end of the cell-cycle. (E) A MATLAB-based dip-detection (Automatic Dip Detection, ADD) method in the mVenus-Hes1 expression profile during the cell-cycle detected the minimum turning point (dip) in the Hes1 expression with 92% accuracy, as compared to manual annotation (FigS6). The duration after the dip (until the second mitosis) showed a tight distribution for all the cells with a mean value of around 11h, while the duration to the dip (from the first mitosis) showed very high variability (6.5-86h) (F). Together, these data showed that Hes1 expression between two consecutive mitoses is biphasic.

We also developed a MATLAB based automatic dip detection (ADD) pipeline (m&m), which detected dips in normalised Hes1 time traces with 92% accuracy (4% false positive and 4% false negative, Fig4E). When mVenus-Hes1 single-cell traces were aligned with the second mitosis as the end-point, the duration of the first episode of Hes1 expression was highly variable, ranging from as short as 6.5h to as long as 72.5h (mean = 18.1h, SD = 11.0h, IQR = 8.1h). By contrast, the position in the time of the dip in Hes1 relative expression in relation to the second mitosis was highly reproducible, which occurred at around 11h after the dip (SD = 3.8h and IQR = 3.2h) (Fig. 4A, D and F).

### 6. The dip in the biphasic Hes1 expression dynamics correlates with the G1-S phase transition of the cell-cycle

Next, we wanted to see how Hes1 dynamics might overlap with the cell-cycle phases. Following on from a previous methodology (30), we developed a lentiviral mCherry-PCNA fusion construct as a single-colour, texture-based reporter of the cell-cycle kinetics. In brief, cells expressing PCNA show smooth nuclear staining in G1 and G2 phases of cell-cycle, while this nuclear staining picks up a punctate pattern during S phase, marking DNA replication foci. The PCNA protein spills over the entire cell during mitoses due to the breakdown of the nuclear envelope (Fig5A and movieS2a and 2b). We performed imaging and analysis on unsorted Cl19 cells virally transduced with mCherry-PCNA so that we could correlate Hes1 dynamics with the cell-cycle kinetics (Fig5A-B). ‘Congo flag’ analysis (Fig5C) showed that in the majority of the cells (93.3%, n = 45), G1 to S transition either happened within the Hes1 dip (5h on either side of the lowest point in Hes1 expression as detected by the ADD pipeline) (50%) or after the onset of the Hes1 dip (43.3%); only a minority of the cells (6.7%) showed G1 to S transition before the appearance of Hes1 dip. The S to G2 transition happened when Hes1 expression started increasing again after the dip in the majority (84%) of the cells (example shown in Fig5B).

**Fig5.**
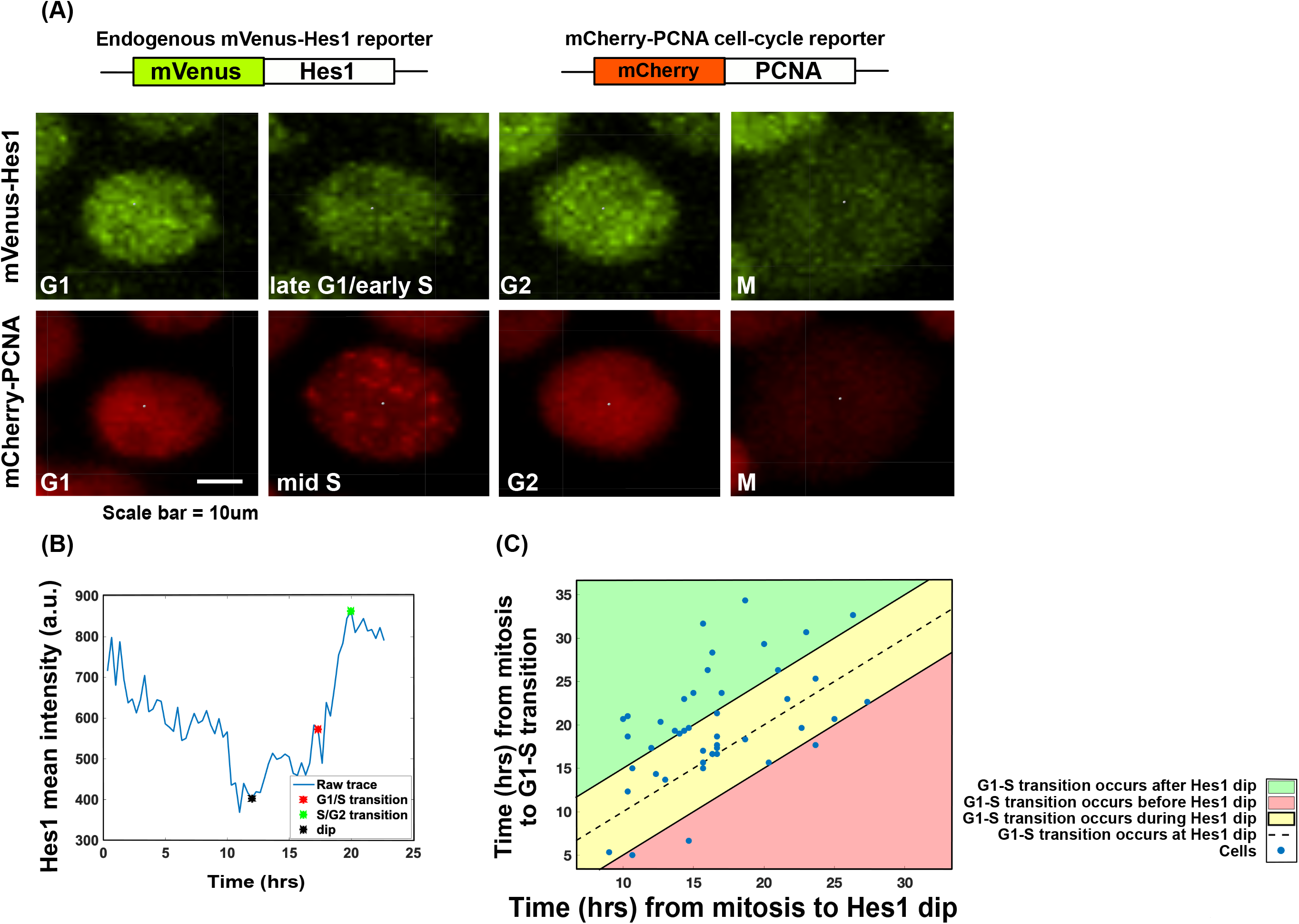
A PNCA-based live reporter shows that the trough in the biphasic Hes1 expression dynamics coincides with the G1-S phase transition of the cell-cycle. (A) Example images of a single cell over time expressing both mVenus-Hes1 and mCherry-PCNA. Cl19 unsorted cells were virally transduced with mCherry-PCNA reporter to show how differentially labelled cell-cycle phases correlate with the Hes1 protein expression dynamics. Scale bar=10um. Schematic of Hes1 and PCNA reporters are shown above the single-cell images. (B) Example of a time trace of Hes1 mean intensity from a single cell expressing PCNA reporter, which was used to mark G1-S and S-G2 transitions, shown as red and green asterisks on top of the Hes1 expression profile. (C) ‘Congo flag’ plots correlating time from first mitosis to G1-S transition (marked by the onset of appearance of PCNA puncti) with time from mitosis to Hes1 dip (with the low expression lasting around 10h as shown in Fig4D) showed that in more than 90% of the cells the G1-S transition occurs either after the dip in the Hes1 expression (green), during the period of the Hes1 dip (yellow) or at minimum turning point (dotted line), with a small minority of cells showing a G1-S transition before the Hes1 dip (pink).

These data from Figs 4 and 5 suggest that the earlier, variable Hes1 peak takes place within the G1 phase of cell-cycle, that the dip in Hes1 expression dynamics mostly precedes the G1 to S transition, and the time-invariant, second Hes1 peak happens between mid-late S and mitosis.

### 7. Gaussian mixture model (GMM) clustering sorts Hes1 dynamics into three distinct classes, with distinct cell-cycle lengths and cellular subpopulation distributions

To understand how Hes1 dynamics interface with the cell-cycle and whether they fall into distinct classes, we ‘stretched’ the Hes1 heatmaps to identical lengths using linear interpolation to represent dynamics at pseudo-time scales. This was followed by classifying these Hes1 pseudo-time series using GMM clustering methods (see m&m and FigS8) to ask whether Hes1 mitosis-to-mitosis dynamics can be classified into discernible, individual groups. Clustering of stretched Hes1 time-series data showed three distinct dynamical Hes1 behaviours during the cell-cycle progression with the following features (Fig6A-B) - Cluster 1 with high-low-high, cluster 2 with medium-low-high and cluster 3 with low-high-low-high relative Hes1 expression. These differences in the dynamics of Hes1 traces during the cell-cycle were also apparent in unstretched but organised data as per above described clusters (FigS9). This along with the analysis of a synthetic control (FigS8) suggested that they are not an artefact of the pseudo-time analysis. The clusters were also evident when the traces were averaged (Fig6B). Further analyses showed that clusters 1 and 2 had similar cell-cycle lengths (mean length = ~25h) and were both enriched in ALDH^High^ bCSCs (Fig6C-D). Interestingly, cluster 3 was enriched in CD44^High^CD24^Low^ bCSCs (Fig6C and D) with highly heterogeneous cell-cycle lengths of up to 90h (mean length 45.2h and IQR 29.5h) (Fig6C). We found that cluster 3 cells share these cell-cycle properties with CD44^High^CD24^Low^ cells as also shown in Fig2B. These data suggest that there are distinct classes of Hes1 dynamics during the cell-cycle and these dynamics are associated with both cellular subtypes and their cell-cycle lengths.

**Fig6.**
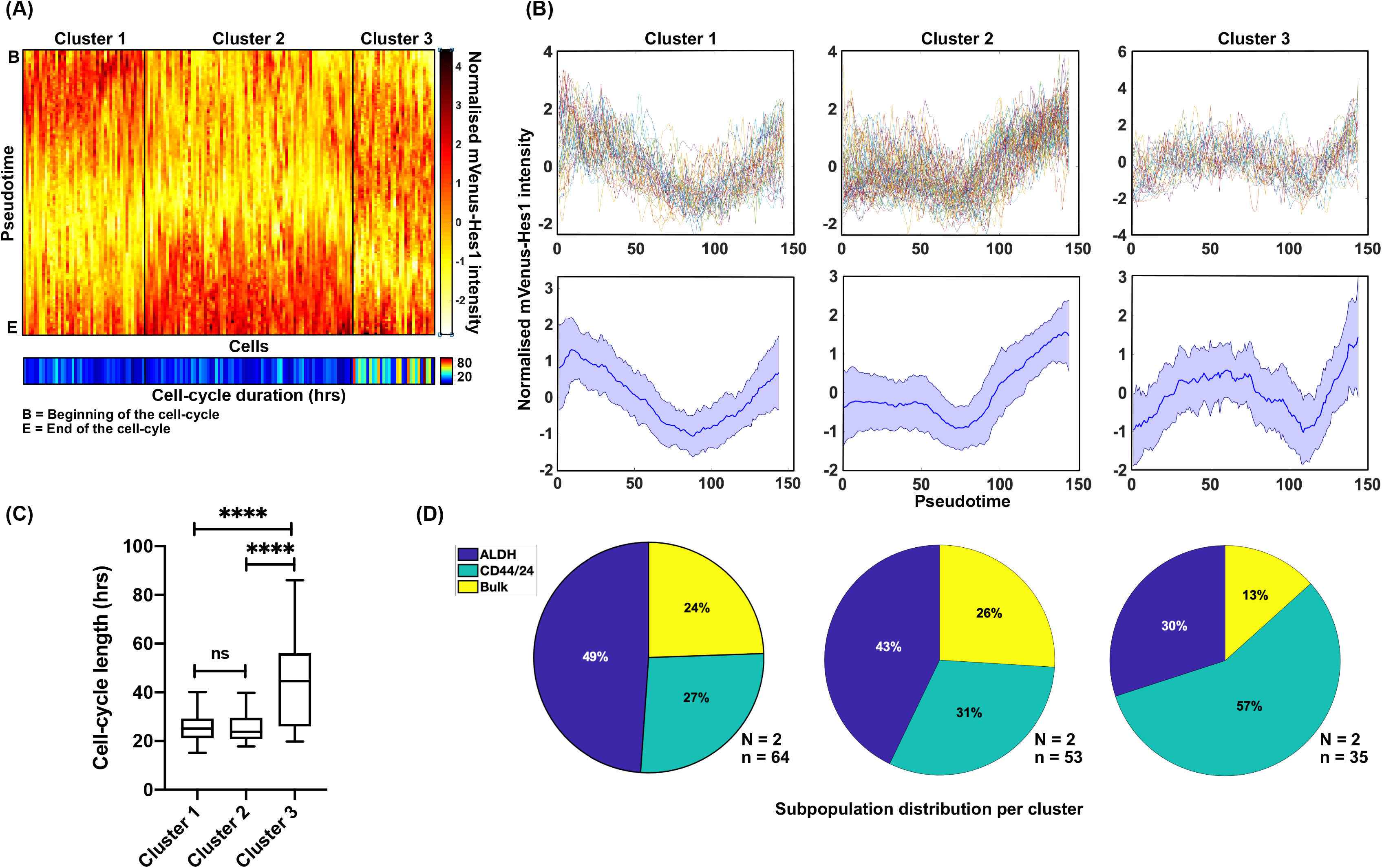
Gaussian mixture model (GMM) shows 3 clusters with distinct Hes1 expression dynamics that map to cellular sub-states and cell-cycle lengths. (A) GMM clustering classifies Hes1 time traces (stretched from mitosis to mitosis, Z scored heat maps) into three distinct classes (clusters 1, 2 and 3, upper panel). The panel underneath shows cell-cycle lengths of individual cells represented as heat maps, showing that cluster 3 is enriched for cells with longer cell-cycles. (B) These clusters can be defined in terms of distinct mean and normalised (Z-scores) Hes1 expression dynamics with cluster 1, 2 and 3 having high-low-high, med-low-high and low-high-low-high Hes1 dynamics, respectively, shown by individual (upper panel) and averaged (lower panel) Hes1 tracks. Dark blue line in the lower panels show the mean Hes1 behaviour per cluster, with lighter blue shading representing the standard deviation errors. (C) Cell-cycle length distributions of individual clusters confirm that cluster 3 consists of cells with the longest and the most heterogeneous cell-cycle, while clusters 1 and 2 are enriched for cells with shorter cell-cycles and less heterogeneity. (D) While cluster 3 is enriched for CD44HighCD24Low bCSCs, clusters 1 and 2 are enriched for ALDHHigh bCSCs. Overall, Hes1 dynamic behaviour can be classified into three classes with distinct cell-cycle lengths and sub-population distributions.

### 8. The onset of the first Hes1 peak correlates with the cell-cycle length

We next sought to understand which feature of the early Hes1 dynamics correlated best with the cell-cycle length. One could hypothesise that cells tend towards a fixed minimum level of Hes1 around G1-S phase transition; therefore, cells with higher peak will take longer to reach this minimum and thus may exhibit a longer cell-cycle (Fig7A). However, we found no evidence for a defined low level of Hes1 that the cells reached as they went through G1 to S transition. Instead, a high starting value of Hes1 correlated with a higher Hes1 value at its lowest level, suggesting a constant fold-change. The fold-change in Hes1 expression between the high Hes1 in G1 and low Hes1 at G1-S had a mean value of 1.6. When we plotted Hes1 intensity at G1 peak against Hes1 intensity at the G1-S trough, a strong positive correlation (R = 0.76) confirmed the invariability in the Hes1 fold-change (Fig7B); these fold-change values did not show a correlation with the cell-cycle length either (R = 0.10, Fig7C). We next looked at the position of the first Hes1 peak by examining the time taken for the cells to reach the first Hes1 peak (Fig7D) and found a good positive correlation between time to peak and the cell-cycle length (R = 0.79) (Fig7E). If we were to remove all values greater than 20h on the y axis, we found that the positive correlation still remained but was slightly weaker (R = 0.55). We also looked at the area under the curve (AUC, Fig7D) for the first Hes1 peak (as a proxy for the shape of the Hes1 peak), and found an excellent positive correlation between the AUC and the cell-cycle length (R = 0.81) (Fig7F), suggesting that the greater amount of time that Hes1 spends above its lower value correlates with longer cell-cycle length. These findings show that the position and the shape of the Hes1 protein peak within the G1 phase are the best predictors of overall cell-cycle length compared to any other dynamic features.

**Fig7.**
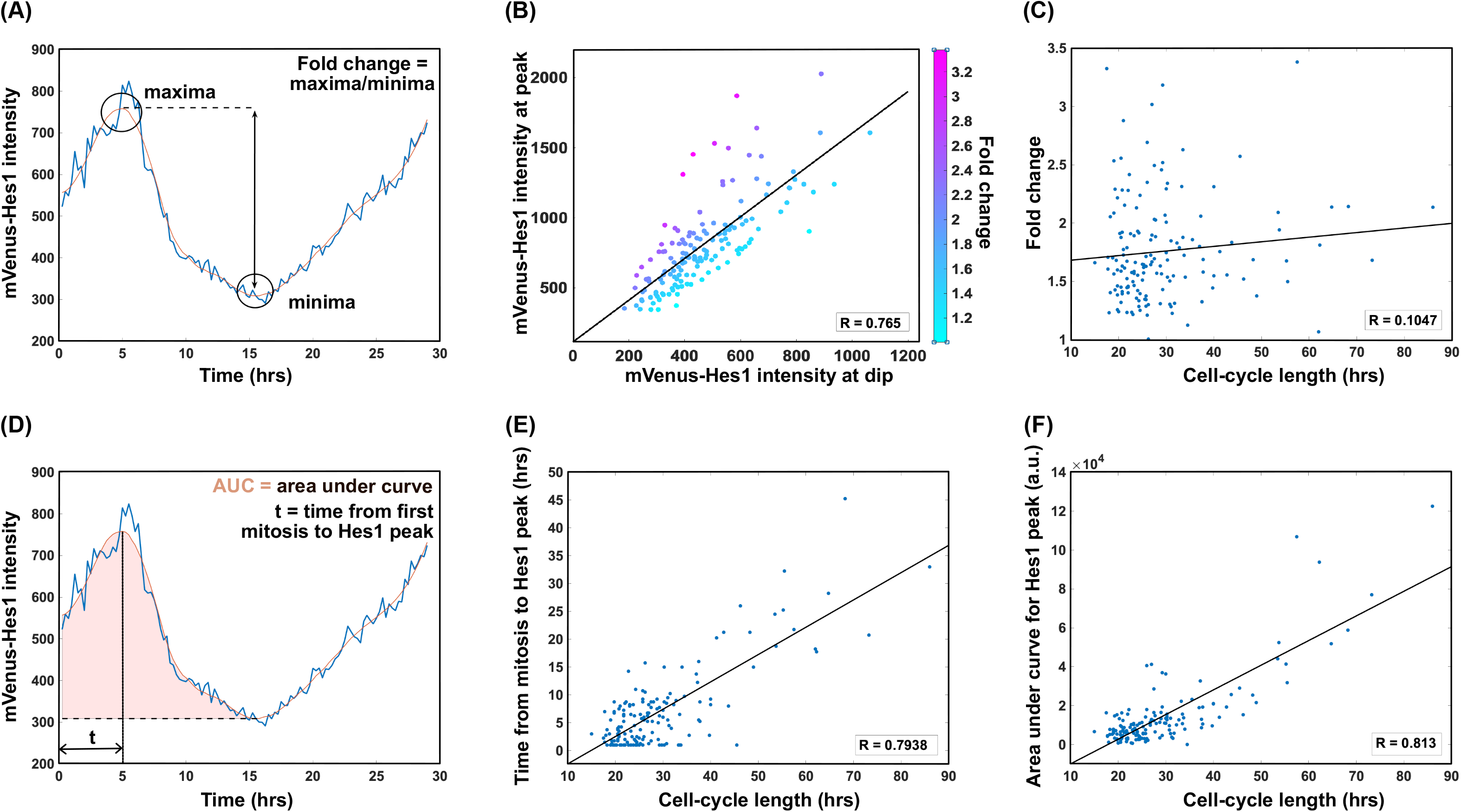
The shape and the onset of the first Hes1 peak can predict the length of the cell-cycle. (A) Example of a Hes1 trace with automatically detected peak of the Hes1 during G1 and the dip at or around G1-S transition. These Hes1 peak (maxima) and dip (minima) values were used to estimate fold-change in Hes1 expression when a cell transitions though the cell-cycle. (B) A strong correlation between Hes1 peak and dip values (R = 0.76) suggested that the Hes1 fold-change values for cells remain within the same range (1.5-2 fold), irrespective of whether a cell started with higher or lower Hes1 intensity. (C) When these estimated fold-change values were plotted against the cell-cycle lengths, no correlation was found (R = 0.10), showing that the cell-cycle lengths in the MCF7 subpopulations were not influenced by the Hes1 fold change values. (D) The same dip detection pipeline was used to estimate how long it took for a cell to reach the peak of Hes1 expression after mitosis (t), and also the area under the curve (AUC) for Hes1 expression during the first phase of the Hes1 expression. (E and F). Both time to peak and AUC showed strong positive correlations with the cell-cycle lengths (R = 0.78 and 0.81, respectively). These data shows that the shape and the onset of the Hes1 expression in its first peak were the best predictors of the cell-cycle lengths in MCF7 subpopulations.

### 9. Mitosis is phase-shifted in relation to the Hes1 oscillator in cells with the longer cell-cycle

The late onset of Hes1 expression in cells with longer cell-cycle prompted us to examine the phase relations between the mitoses at the beginning of a cell-cycle and the Hes1 periodic wave. We constructed phase relation diagrams using Hilbert Transform of the dynamic Hes1 traces using the MATLAB platforms (Fig8A). Phase diagrams for cells (histograms in Fig8B and circular plots in Fig8C) suggested that for the majority of cells from clusters 1 and 2 with shorter cell-cycles, the mitoses are distributed around the Hes1 peak and the descending slope towards the Hes1 trough. Since the majority of the cells in the culture divide around the peak (0 and 2π, Fig. 8Β, first panel) we defined this as the dominant phase (“in-phase”) of the Hes1 oscillator and mitosis. Interestingly, cluster 3 cells originate from divisions that preferentially take place at or around a trough of the Hes1 wave (Fig. 8B, second panel), therefore, these cells start their cell-cycle with a relatively low Hes1, which ascends to an apparently delayed peak of Hes1 expression in G1; we termed this as “out-of-phase” or “phase-shifted” register of the Hes1 oscillator with mitosis (Fig8C; also see FigS10A). Furthermore, when we presented our data for cellular subpopulations and additionally splitting it into cells with cell-cycle length up to 40h and cells with cell-cycle longer than 40h (Fig8D), it was clear that almost three quarter of the cells with longer cell-cycle lengths (>40h) originated from the mitoses which were out of phase with the Hes1 wave and that such cells were preferentially of CD44^High^CD24^Low^ type (Fig8D). Taken together, these data show that majority of the MCF7 population exhibit in-phase mitoses around Hes1 peak, resulting in shorter cell-cycles. In contrast, out-of-phase mitoses, i.e. mitoses occurring closer to a retrospective trough (i.e. one that is only apparent when the next cell division is considered) result in cells with longer cell-cycle, which were mainly representing the CD44^High^CD24^Low^ type cancer stem cells (Fig8C-D, FigS10A and S11D, right hand side example).

**Fig8.**
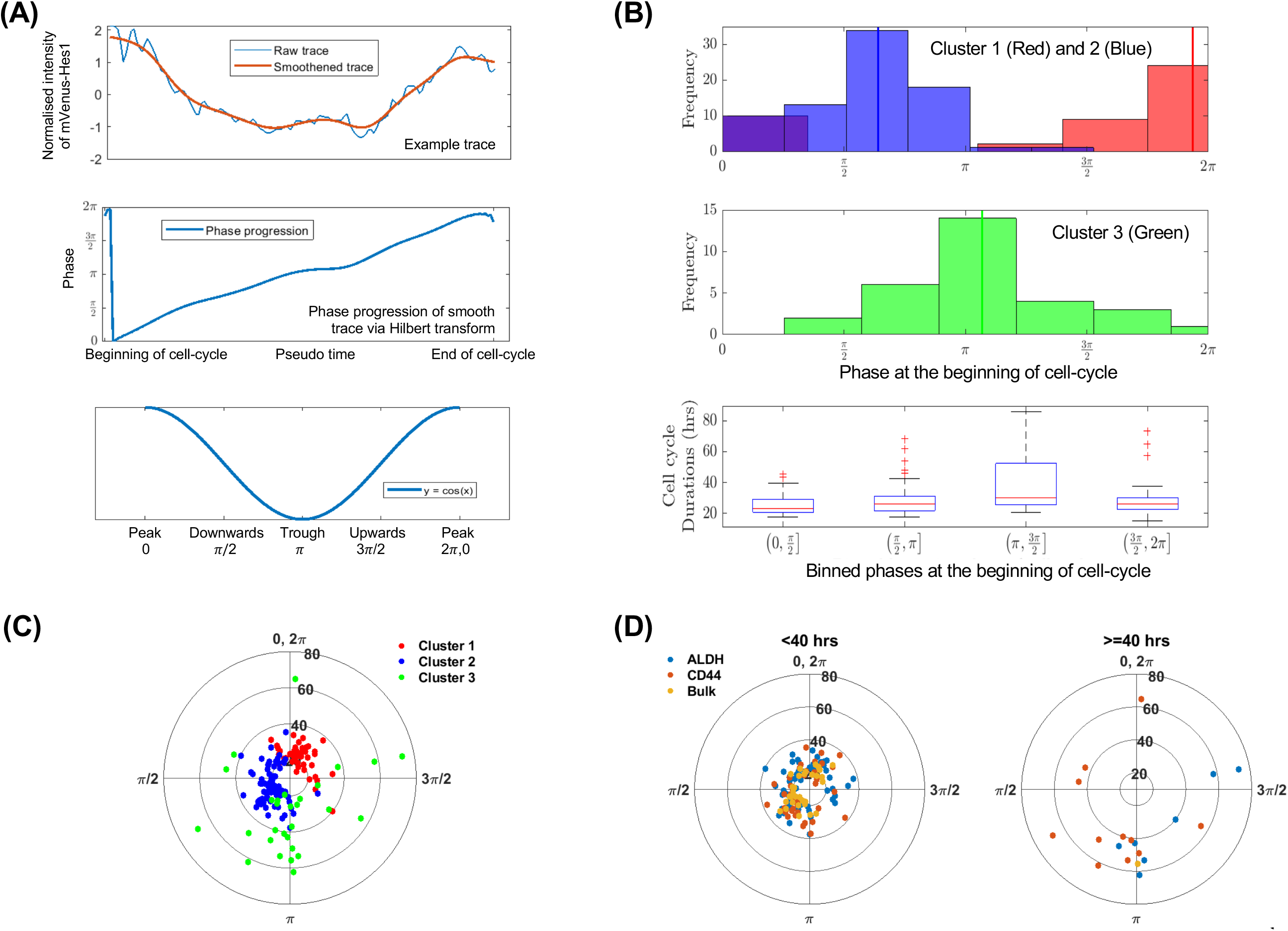
The ‘phase register’ of mitoses with the Hes1 oscillator and its influence on the length of the cell-cycle - mitosis is phase-shifted in relation to the Hes1 oscillator in cells with the longer cell-cycle. (A) Upper panel shows an example of a normalised (z-scored) Hes1 pseudo-time series (blue) over one cell cycle (mitosis to mitosis) with smoothened trace (orange). Middle and lower panels show phase reconstruction for the smoothened trace (above) via Hilbert transform with phase readouts of an example cosine wave on x-axis in the lower panel. Phases 0 or 2π represent peak and phase π represents a trough in the Hes1 wave in one cell cycle. (B) Histograms showing where on the Hes1 wave, starting mitoses took place for cells from clusters 1 and 2 (upper panel) and cluster 3 (middle panel) using the bottom panel in (A) as a reference. For cells from clusters 1 and 2, the majority of the mitoses took place around the descending or at the Hes1 peak (upper panel). For cells from cluster 3, the majority of the mitoses happened at an apparent Hes1 trough. These cells show a delayed Hes1 peak in G1 which makes the starting level of Hes1 at the exit of the previous mitosis appears retrospectively as a trough. (C) Scatter plot in polar coordinates where the angle (reading anti-clockwise from the vertical) represents the phase read-out at the mitosis point preceding a cell-cycle. The distance from the centre of the circle represents the cell-cycle length of that cell. Cluster 1 cells are in red, cluster 2 in blue and cluster 3 in green. (D) Scatter plots similar plot to (C), except here ALDH^High^ bCSCs are in blue, CD44^High^CD24^Low^ bCSCs are in orange and Bulk cells are in yellow. The left panel shows cells with cell-cycle lengths up to 40h and the right panel shows cells with cell-cycle lengths over 40h. Note that cluster 3 and CD44^High^CD24^Low^ cells particularly the ones that have long cell cycles, preferentially occupy the lower half of the scatter plot (between π/2 and 3π/2 which suggests that the mitoses that give rise to these cells preferentially take place at or around a trough in the Hes1 wave. Overall, histograms in B and Scatter plots in C and D show that preceding mitoses for cells with longer cell-cycles (which mostly belong to cluster three and enriched for CD44^High^CD24^Low^ bCSCs), register with the Hes1 wave ‘out of phase’, i.e. the mitoses occur around a trough, while cells with shorter cell-cycles are more likely to divide around a peak of the Hes1 wave.

### 10. Cancer stem cell subpopulations differ in their switching probabilities

Cancer stem cell plasticity that underlies non-genetic heterogeneity postulates that cells are not fixed in their properties over time. Therefore, we looked for evidence of switching behaviour between clusters 1, 2 and 3, defined by the Hes1 dynamics over 2 generations (normalised and clustered separately; Fig9A and FigS10). We found that cells do indeed switch between cluster classifications and these ‘cluster transitions’ are seemingly random i.e. cells from any cluster in generation 1 (mother cell) can fall into any cluster classification upon division into generation 2 (daughter cell) (Fig9A). However, they are weighted in that some transitions are more likely than others (e.g. 3 into 1 is more likely than 3 into 3). Transition into cluster 3 (out of phase behaviour) is less frequent in all clusters than transitions into clusters 1 and 2. Moreover, cells from cluster 3 switches to cluster 1 and 2 very often (i.e. cells with “out-of-phase” behaviour almost invariably divide back into phase), than clusters 1 and 2 that tend to stay in clusters 1 and 2 (Fig9B). We also found that cells that have been sorted for CD44^High^CD24^Low^ cells switch to or from cluster 3 more often than ALDH^High^ cells. This is consistent with the cluster switching frequencies above, since cluster 3 is enriched in CD44^High^CD24^Low^ cells, and CD44^High^CD24^Low^ cells tend to fall into cluster 3 more often than ALDH^high^cells (Fig9C). In all cases, transitions into or out of cluster 3 was accompanied by an elongation or a shortening of the cell-cycle length and Hes1 periodicity (FigS10 and S11B), which further strengthened the link of the phase register with the cell-cycle length.

**Fig9.**
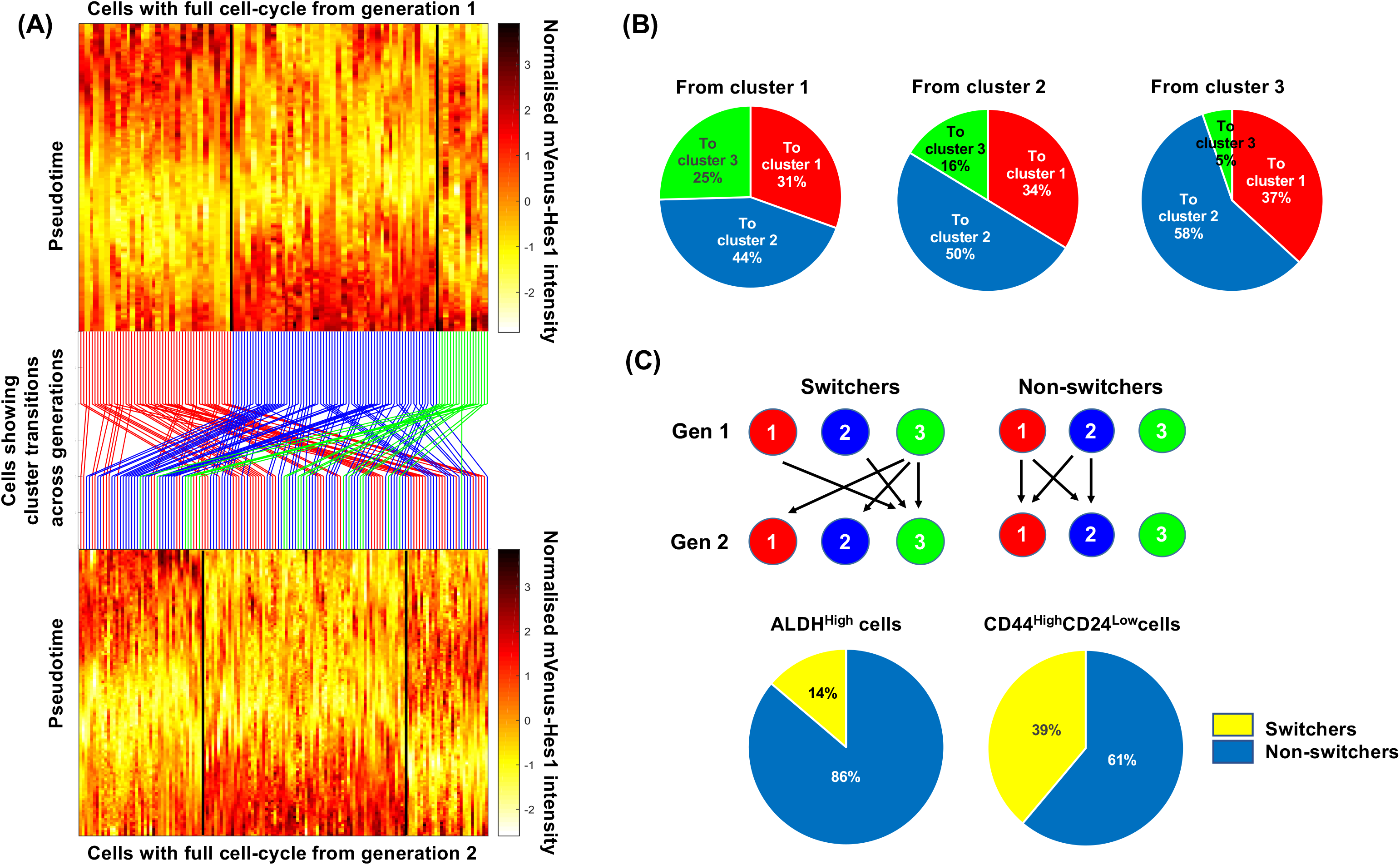
Cells exhibit weighted cluster transitions, with CD44^High^CD24^Low^ cells showing higher switching behaviours, compared to the ALDH^High^ cells. (A) Heatmaps of normalised (z-scored) Hes1 pseudo-time series with full cell-cycle classified as clusters in two consecutive generations (gen 1 on top and gen 2 at bottom). Cells across these two generations have been connected through lines (red, blue, green, corresponding to clusters 1, 2 and 3 as read from left to right) in the middle of these two heatmaps, showing how individual cells jump across clusters when they go from one to another generation. (B) Cluster transition rates show that these cluster transitions were not completely random, but weighted, with cluster 3 cells more likely to transition out into clusters 1 and 2 (right side pie chart), but clusters 1/2 were more likely to stay within clusters 1/2, and less likely to transition into cluster 3 (left and middle pie charts). (C) Upper panel defines switchers as cells moving in or out of cluster 3, while non-switchers as cells staying within clusters 1/2, irrespective of their cell generations. CD44^High^CD24^Low^ cells are more likely to switch (39%) as compared to ALDH^High^ cells (only 14%) (lower panel), corroborating the previous findings that CD44^High^CD24^Low^ cells are more heterogenous in cell-cycle lengths.

### 11. Bidirectional functional interaction of the Hes1 oscillator and the cell cycle

To ask whether the influence from Hes1 oscillations to cell-cycle length is causative, we directly manipulated the Hes1 expression dynamics. We made use of a viral expression system wherein mVenus-Hes1(cDNA) is expressed under the Ubc promoter (Ubc:mVenus-Hes1) for a sustained rather than a dynamic mVenus-Hes1 expression in the parental MCF7 cells (Fig10A-F). MCF7 cells expressing this reporter showed variability in the levels of the exogenous mVenus-Hes1 expression. To avoid any impact asserted by the over-expression of mVenus-Hes1, we focused on cells with low to medium expression levels of the exogenous mVenus-Hes1. Periodicity analysis of mVenus-Hes1 traces using the LSP platform and the heat maps generated from the normalised expression confirmed that the mVenus-Hes1 expression generated from this viral reporter was not periodic (Fig10A-B). Any protein fluctuations were aperiodic and the characteristic Hes1 dip marking G1 to S transition at a fixed position before mitosis was absent (Fig10C-D). When these low-medium Ubc: mVenus-Hes1 expressers were compared to the control MCF-7 cells expressing nuclear Venus alone (Ubc:NuVenus), the experimental cells showed significantly longer cell-cycle lengths (Fig10E) and overall, fewer cell divisions (Fig10F). Sustained expression of Hes1 did not only elongate the cell cycle but this was also accompanied by an increase in the proportion of CD44^High^CD24^Low^ cells (FigS12); this may happen as a result of sustained Hes1 expression suppressing the switching behaviour of CD44^High^CD24^Low^ cells. Thus,sustained Hes1 expression under a ubiquitous promoter alters cell-cycle length and cell fate in MCF-7 cells. Together, these data suggest that dynamics of Hes1 expression is required for smooth progression of the cell-cycle and cell fate specification.

**Fig10.**
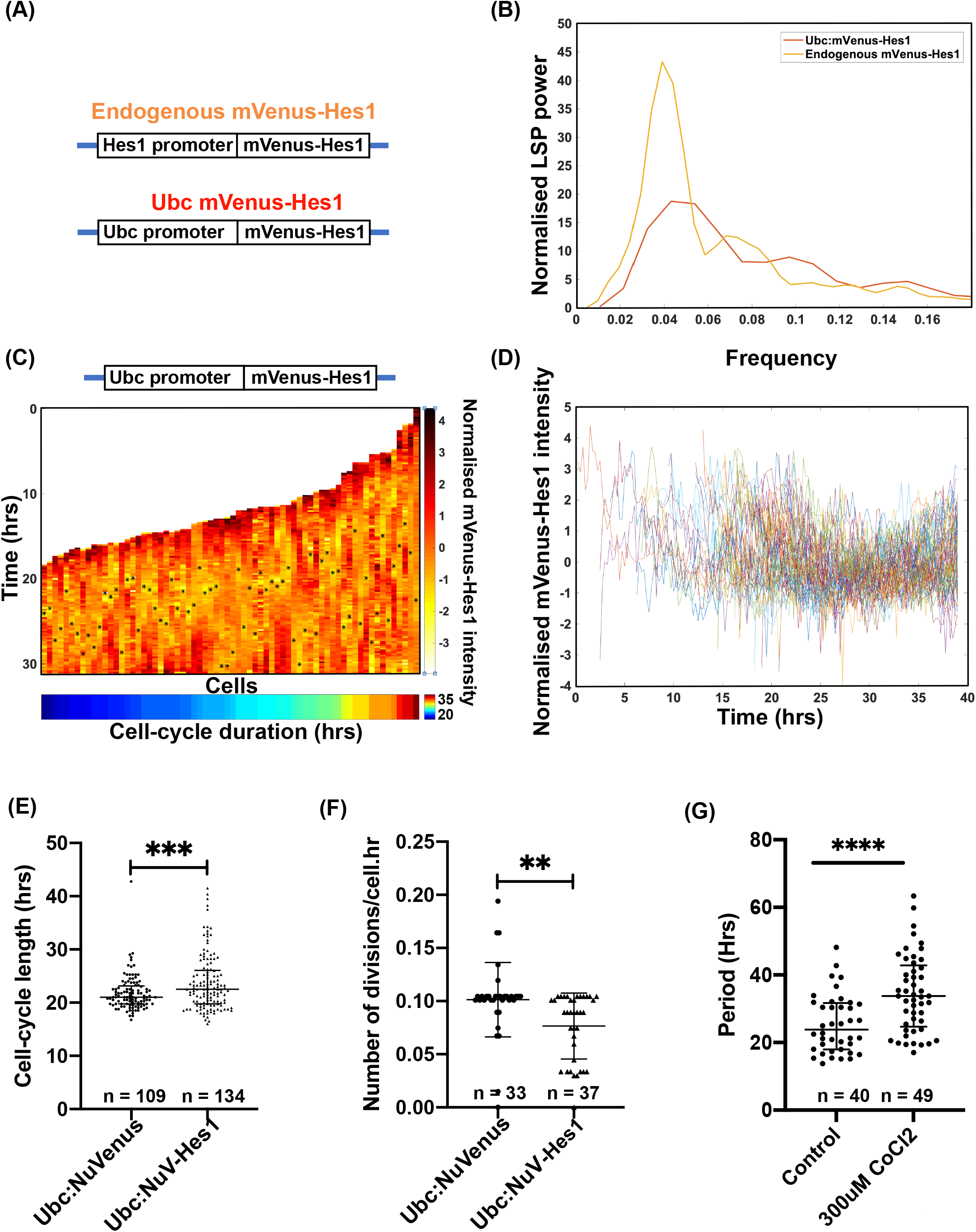
Bidirectional interaction of Hes1 oscillations and the cell cycle. **(A-F) Dampening Hes1 oscillations by sustained expression of Hes1 elongates or impairs the cell-cycle.** (A) Schematic of constructs for endogenous expression of mVenus-Hes1 (orange) and mVenus-Hes1 under the Ubc promoter (red). (B) Normalised (scaled between 0-100) LSP power shows a strong peak in the power spectrum obtained from the endogenous mVenus Hes1 expression, in contrast to the mVenus-Hes1 expression under the Ubc promoter. This suggested that endogenous mVenus-Hes1 shows coherent oscillations while the same protein does not oscillate over time when expressed under the Ubc promoter. (C) Traces of Ubc-mVenus-Hes1 cells, normalised and arranged as a heatmap from mitosis to mitosis with time running vertically from top to bottom. These are aligned in time at the second mitosis of each cell. The time traces have been arranged in increasing order of the cell-cycle length. The dip in expression found by our ADD pipeline (black asterisk) shows random dips unlike in the Endogenous mVenus-Hes1 time traces (Fig4A and D). (D) Ubc-mVenus-Hes1 time tracks from cells arranged as normalised time series traces from mitosis to mitosis, aligned in time at the ending mitosis, confirming the absence of a common dip in expression before division, compared to the endogenous mVenus-Hes1 as shown in (Fig4A and D). When Ubc:mVenus-Hes1 cells were compared against control cells expressing Nu-Venus alone for their cell-cycle properties, experimental cells were found to exhibit longer cell-cycle (E) and to have lower number of cell-divisions (F). Together these data show that Hes1 oscillations are required for the smooth progression of the cell cycle. **(G) Blocking cell-cycle using CoCl2, a hypoxia mimetic, elongates the periodicity of Hes1.** This elongated Hes1 represents the period of Hes1 in the absence of an input from the cell-cycle, possibly representing the ‘free running’ period when Hes1 oscillations and the cell-cycle are not coupled. Control n=40 cells, 300uM CoCl2 n=49 cells.

In the experiments above, to exclude the possibility that any effect of Ubc:Hes1 on cell-cycle and cell fate is due to Hes1 overexpression above its physiological level, we transduced M18 cells (CRISPR/Cas9 line expressing mScarlett-Hes1, see FigS4A-C and m&m) with lentivirus expressing Ubc:mVenus-Hes1. We then compared the overall Hes1 fluorescence intensity in transduced stable lines with that of endogenously tagged M18 or C19 cells (FigS13A). We found that the overall Hes1 intensity in transduced cells was very similar to mScarlet-Hes1 or mVenus-Hes1 intensity in untransduced M18 or Cl19 cells respectively. (FigS13B). The collected mean intensities of mVenus and mScarlet were level-matched against each other for these analyses, as shown in supplementary figs 13B.

To understand why the addition of Ubc:mVenus Hes1 expression does not raise the total Hes1 level, we looked at its impact on the endogenous Hes1 expression (tagged with mScarlet) and we found an almost complete suppression of the endogenous expression (FigS13A and C). We suggest that this is due to the property of autorepression of Hes1 gene on its own promoter, which results in Ubc:mVenus Hes1 expression repressing the endogenous gene but not the other way around. We have made similar observations in mouse Hes1 in the context of neural stem cells (31). In conclusion, the suppression of endogenous Hes1 gene expression by exogenous mVenus-Hes1 driven off the Ubc promoter serves to maintain the total Hes1 at a physiological mean level thus, linking the phenotypic effects to altered dynamics rather than an altered mean level of expression.

Furthermore, to test for functional evidence of the reciprocal interaction between the cell-cycle and the Hes1 oscillations, we inhibited cell-cycle progression by treating cells with CoCl_2_ (see m&m) (32) and looked at Hes1 oscillations. We found that Hes1 oscillations continued even after the cell-cycle was blocked but the periodicity was longer (Fig10G). These data suggest that the cell-cycle and the Hes1 oscillator represent oscillations that can proceed independently with a longer “free running” period, but are normally coupled.

### 12. Sustained Hes1 expression impacts the dynamics of the cell-cycle regulators

To understand and strengthen the functional link between the Hes1 dynamics and the cell-cycle, we made use of the mVenus-p21 CRISPR MCF7 line generated using a similar CRISPR tagging strategy (33). Using single-cell, live-imaging approach, we monitored the expression of p21 under conditions of control and sustained Hes1 expression. We found that under control conditions, p21 had a peak of expression in the G1 phase, prior to the G1-S transition, as previously reported (33). But when we transfected the mVenus-p21 cells with a plasmid expressing sustained mScarlet-Hes1 under the Ubc promoter, p21 also lost its expression peak, showed sustained expression and cells did not divide (FigS14). This data suggested that Hes1 may interact with the cell-cycle machinery via its regulation of the dynamics of the key cell-cycle molecules such as p21.

## Discussion

The oscillatory protein expression of Hes1, a transcriptional repressor of the helix-loop-helix family, has been shown to be important for cell-state transitions in neural progenitor cells (8) (9) (21). Even though such progenitor cells divide, it was an open question how Hes1 oscillations relate to the process of cell division. In this paper, we have asked whether Hes1 oscillations are observed in breast cancer cells and whether they have a reproducible and functional relation to their cell-cycle.

Using an endogenous CRISPR/Cas9 fluorescent fusion reporter, we show that Hes1 oscillates with an average periodicity of 25h in MCF7 breast cancer cells. This is significantly longer than the previously reported (2-3hr) ultradian periodicity in mouse cells (31 and references within). We did notice that similar ultradian oscillations are also “embedded” in the longer duration oscillations we report in this paper (as is apparent in the example Hes1 time trace shown in Fig4D); they have a periodicity of 5-6 hours possibly due to the slower pace of oscillation in human cells, as reported for somitogenesis (20) (34). In this study, continuous live-imaging of 4-5 days duration, which far exceeded any previous live-imaging for Hes1 protein using mouse-based (or any other) experimental systems, uncovered these novel, non-ultradian, circadian-like oscillations of Hes1 that we focused on, as they showed clear correlations with both cell-cycle and cell fate, as compared to the ultradian oscillations (FigS6).

MCF-7 cells are a good model to study non-genetic heterogeneity in breast cancer because they contain “bulk’’ cells, i.e. cells with limited potential for tumour (re-)initiation, as well as populations of cells that can act as cancer stem cells when transplanted *in vivo*, namely ALDH^High^ epithelial-like stem cells and CD44^High^CD24^Low^ mesenchymal-like stem cells (18). Here, we report that the majority of the MCF7 cells (represented by bulk cells) have a consistent cell-cycle length of approx. 24h while cells sorted for known cancer stem cell markers show greater variability in their cell-cycle length. Interestingly, stem cells expressing CD44^High^CD24^Low^ markers, which are more likely to become dormant *in vivo*, show more significant variability than the average of the population and significantly longer cell-cycles, which may predispose them to quiescence, in the right environment.

To understand how these long cell-cycles may be generated, we asked whether the cell-cycle interfaces with the Hes1 oscillations in a manner which may be causal. Indeed, we report here that, in most cells, Hes1 oscillations have a reproducible relationship to the cell-cycle such that mitosis tends to take place at, or around, the peak of Hes1 expression (‘In-phase’ register). Following mitosis, a trough of Hes1 expression precedes or coincides with the G1 to S transition. After this trough, Hes1 protein concentration picks up again as the cells progress to the second mitosis. Thus, because of the tendency of mitosis to take place around the peak, Hes1 expression appears biphasic within every cell-cycle, with two periods of increased protein concentration in G1 and G2. This biphasic expression may reconcile conflicting activities of Hes1, which is known to repress both activators (such as CyD, CyE and E2F) (14) (15) (16) and inhibitors of the cell-cycle (such as the CDKIs p21 and p27) (12) (13), by providing a temporal separation of repressing phases with a period of de-repression at the trough. In fact, most of these proteins show dynamic expression during the cell-cycle (33) (35) and we have shown that the expression of p21 is no longer dynamic when Hes1 is sustained, suggesting that Hes1 expression dynamics normally contributes in shaping or “sculpting” the dynamics of the cell-cycle molecules. Future experiments will show whether de-repression of molecules such as E2-F1 or CyclinE (a target of Hes1 in MCF-7 cells and known for its function in G1-S transition, https://www.encodeproject.org/experiments/ENCSR109ODF/) may be due to the dip in Hes1 expression and in turn may time the G1-S transition. The trough in Hes1 expression at around S-phase is most likely transcriptional as it is not observed when Hes1 is expressed under a ubiquitous promoter and may be due to the loss of Notch signalling in S-phase, reported in developmental systems (36).

The observed variability in the cell-cycle length was specifically associated with the first peak of Hes1 expression during the cell cycle, while the time between the Hes1 trough in G1/S and the subsequent mitosis was less variable. This fits well with reports in the literature wherein neural stem cells, the cell-cycle length is primarily determined by the length of G1 (37). Given this, we further analysed the expression of Hes1 in G1 phase of cell-cycle, to see which Hes1 expression feature shows the best correlation with the cell-cycle length, implying a functional decoding role. One could hypothesise that cells that start with high Hes1 level would take longer to reach a set low Hes1 level, which could be required for G1-S. However, we found that neither the starting levels nor the mean Hes1 levels during the cell-cycle showed a good correlation with cell-cycle length. Furthermore, the trough of Hes1 levels had a little varied fold-change in relation to the starting level, suggesting that there is no fixed absolute level of low Hes1 that the dynamics tend to. This finding implies that any downstream targets of Hes1 sense a fold-difference rather than an absolute minimum level of Hes1 at or around G1-S transition during the cell-cycle progression. Thus, Hes1 oscillations may be an underappreciated example of a Fold Change Detection (FCD) system, a possible feature of gene regulatory networks with feedback (38).

Instead, we found that the position (onset) and/or the “area under the curve” (shape) of the first Hes1 expression peak, as recorded in live-imaging traces, was a better predictor of cell-cycle length than its levels of expression. Cells with a delayed onset Hes1 peak during the G1 phase of cell-cycle, that is, a Hes1 peak positioned with some temporal separation from the previous mitosis, tended to have a longer cell-cycle going forward. Such a delayed Hes1 peak in G1 supersedes the previous G2 peak which is no longer a peak but part of a new, ascending Hes1 trend (picture abstract and example trace in Fig3D-F). Therefore, these cells are now “out of phase” with the cell-cycle and the Hes1 period is elongated when one measures the distance from the delayed peak to the Hes1 peak in the preceding G1 (Fig S11B). Indeed, phase reconstruction showed that cells with longer cell-cycles tended to have an unusual phase register with the cell-cycle such that their mitoses tend to take place at around Hes1 trough rather than at the peak of Hes1 wave. This delayed Hes1 expression peak in G1 appears to have a knock-on effect in also delaying the trough, which if Hes1 has a gating effect, would delay the G1 to S transition and the overall cell-cycle length of these cells (FigS10B).

Furthermore, CD44^High^CD24^Low^ stem-like cells were more likely than ALDH^High^ stem-like cells to show a late-onset Hes1 peak in G1 and thus, a phase-shifted registration of Hes1 dynamics with the cell-cycle. However, the phase registration was not set in time and cells belonging in different cluster classifications (with respect to their alignment of Hes1 dynamics with the cell-cycle), switched between behaviours randomly but with weighted probabilities. CD44^High^CD24^Low^ cells showed the highest probability of switching Hes1 dynamics in and out of phase with the cell-cycle, which fits well with the greater heterogeneity in cell-cycle length in these cells. Future experiments to understand the basis of the increased switching behaviour of these cells will be important, as this property may underlie the higher propensity of these cells to become dormant *in vivo* (18).

To investigate further if Hes1 oscillations and the cell-cycle represent coupled systems, we mis-expressed mVenus-Hes1 under a ubiquitous promoter (Ubc). In cells expressing this construct, Hes1 did not oscillate showing only random fluctuations in its expression. Since previous finding showed that under conditions of overexpression, high Hes1 levels are associated with cellular dormancy (22) (39) (40), we have taken care to show that the level of Hes1 is comparable to physiological levels. In this population, we found a higher proportion of CD44^High^CD24^Low^ cells and a higher proportion of cells with a longer cell-cycle or that failed to divide, suggesting that Hes1 oscillations are necessary for an efficient cell-cycle with a consistent length. Reciprocally, blocking cell division completely with CoCl_2_, a hypoxia mimetic known for blocking cell-cycle and pushing cells into dormancy (32), elongated the Hes1 period and reduced their amplitude. These results suggest that the cell-cycle and Hes1 oscillations represent periodic systems which can proceed independently but are normally reciprocally coupled, with a preferred, fixed phase relationship.

In conclusion, Hes1 expression appears heterogeneous in snapshots of ER+ breast cancer cells but this is due to asynchronous temporal oscillations with a circadian-like periodicity. Our findings suggest that most likely through regulation of cell-cycle molecules, these Hes1 dynamics sculpt the cell-cycle to an efficient nearly 24h periodicity. Under normal conditions, it appears that cell-cycle and Hes1 oscillations are coupled systems and their dynamic phase registration is important in controlling the cell-cycle length and cell fate. In that respect, our findings have some analogies to developmental systems, where the phase shift of periodic signalling molecule encodes information for cell fate (34). Notably, a particular sub-type of stem-like cancer cell, the CD44^High^CD24^Low^ cells, show a higher rate of in- and out-of-phase switching and greater heterogeneity of cell-cycle than ALDH^High^ cancer stem-like cells. This suggests that frequent misalignment of the Hes1 oscillations with the cell-cycle may eventually contribute to the known tendency of CD44^High^CD24^Low^ cells to drop-out of the cell-cycle *in vivo*, which may have important implications for the progression and relapse of the breast disease. We suggest than an important area of future work is the interface of Hes1 with the circadian clock, which also shows coupling with the cell-cycle (41) and has important implications for the chronobiology of cancer (42) (43).

## Materials and methods

Please refer to the supplementary materials section for all the details concerning materials and methods.

## Acknowledgements and funding sources

We thank Profs Robert Clarke from the Manchester Breast Centre and Keith Brennan from the Cell Matrix Centre, FBMH, UoM for providing the parental MCF7 cells and the Human Hes1 cDNA plasmids, respectively. We are grateful to Dr Gareth Howell and Mr Michael Jackson from the FACS facility, FBMH for all their help and expertise in sorting and analysing the cell lines used in this study. We thank Prof Galit Lahav (Harvard Medical School) and Dr Jacob Stewart-Ornstein (University of Pittsburgh) for their prompt responses and for making available their MCF-7 p21 CRIPSR line. We also thank the Genomics facility, FBMH for their sequencing services and Dr Anthony Adamson, from the Genome Editing Unit, FBMH for his guidance on the CRISPR/Cas9 designing and implementation. The work was sponsored by the Wellcome Trust grant no 106185/Z/14/Z to Prof Nancy Papalopulu.

