## Supplementary material for "Differential phase register of Hes1 oscillations with mitoses underlies cell-cycle heterogeneity in ER+ breast cancer cells": Supp Info

PNAS
www.pnas.org

#### Supplementary Information for

**Differential phase register of Hes1 oscillations with mitoses underlies cell-cycle heterogeneity in ER+ breast cancer cells.**

Nitin Sabherwal, Andrew Rowntree, Elli Marinopoulou, Tom Pettini, Sean Hourihane, Riba Thomas, Jochen Kursawe, Ximena Soto and Nancy Papalopulu

Correspondence –
Nancy Papalopulu and Nitin Sabherwal


**This PDF file includes:**

Materials and methods
Legends for supplementary figures
Legends for Movies S1 to S2
SI References
Figure S0 - Abstract in picture
Figures S1 to S14

**Other supplementary materials for this manuscript include the following:**

Supplementary movies 1, 2a and 2b

#### Materials and methods

##### Cell culture

Parental MCF7 and any other reporter lines derived from it were propagated in high Glucose DMEM (Sigma-Aldrich, D6429) supplemented with 10% fetal bovine serum (FBS, Sigma-Aldrich) and 1% Penicillin-Streptomycin (Pen-Strep, Sigma-Aldrich), at $37^0$C under 20% O2 and 5% CO2 culture conditions. Adherent cells were passaged twice weekly using 1X Trypsin-EDTA solution (Sigma-Aldrich) to dissociate them from the culture flask. Cells were routinely checked for any mycoplasma contamination using in-house established PCR based testing method.

##### Plasmids and viral reporter lines

To generate pCS2-HA-Hes1 plasmid (expressing human Hes1 fused to HA epitope under CMV promoter), Hes1 was PCR-amplified from the plasmid pBS-SK-HuHes1 using primers with restriction sites introduced. Purified and digested PCR product was sub-cloned in the pCS2-2HA plasmid. Clones were sequence-verified before using them to transfect MCF7 cells using Lipofectamine 3000 using the manufacturer's protocols (Life Technologies).

To generate III generation viral plasmid constructs, we used pLNT-Ubc:mVenus plasmid (Ubc driving mVenus expression) (1) as the backbone for any further subclonings. To generate pLNT-Ubc:NuVenus (nuclear mVenus driven by Ubc promoter) plasmid, mVenus cDNA was amplified using the forward primer containing SV40 NLS and the reverse primer. Digested PCR product was subcloned back in the same vector backbone, and selected clones were sequence verified. Similarly, for the plasmid pLNT-Ubc:mCherry-PCNA (mCherry-PCNA driven by Ubc promoter), primers with introduced restriction sites were designed to amplify mCherry cDNA and human PCNA cDNA separately. Both PCR products were digested and subcloned in the original pLNT vector one after the other, followed by sequence verification of the selected clones. Plasmid for expressing mVenus fused to human Hes1 cDNA under the Ubc promoter (pLV-Ubc:mVenus-Hes1) was obtained commercially from VectorBuilder. Details for all primers and cloning strategies are available on request. Lentivirus production and transduction of MCF7 cells were performed as described before (1). To maximise the cell population expressing fluorescent reporter of interest, transduced cells were FACS sorted on BD Influx Cell Sorter (BD Biosciences, UK).

##### N-terminal in-frame fluorescent tagging of human endogenous Hes1 gene in MCF7 cells using CRISPR-Cas9 genome editing technology

The donor plasmid for N-terminal tagging of endogenous human Hes1 DNA in MCF7 cells with fluorescent monomeric Venus (mVenus) tag was commercially synthesised from GenScript. It consisted of 800 bps upstream of the Hes1 start codon (5' homology arm/5' HA), 800 bps downstream of the Hes1 start codon (3' HA) and in between these two HAs, a mVenus cassette consisting of ATG-3XFLAG-mVenus(cDNA)-Linker-HA, all cloned in a pUC57 based vector backbone (Fig1A). In the donor plasmid, silent mutations were introduced near the PAM sites where the Sp-Cas9 protein would cut the Hes1 genomic DNA. CRISPR-Cas9 machinery inserts the mVenus cassette downstream of the endogenous Hes1 ATG through homology-directed repair (HDR) reaction using donor plasmid as a template (Fig 1A). gRNAs against the human Hes1 were designed using the Wellcome Trust Sanger Institute portal (https://wge.stemcell.sanger.ac.uk//). The gRNA 5'-gaaaaattcctcgtccccggtgg3-' was commercially obtained from Integrated DNA Technologies (IDT) and used in the CRISPR-Cas9 reaction. tRNA and purified Sp-Cas9 proteins were also obtained from IDT. gRNA (200µM) and tRNA (200µM) stock solutions were mixed in IDT duplex buffer, so their final concentrations were 48µM. The mixture was heated at 95°C for 5 min, followed by slow cooling at room temperature. To make the RNP complex, 5µL of gRNA:tRNA complex was mixed with 2µL of Cas9 (61µM) and 4µL of IDT duplex buffer, making the final concentrations of both gRNA and tRNA as 20.4µM and Cas9 as 11.09µM. The RNP complex was incubated at room temperature for 10 min before it was mixed with 2µg of donor plasmid. The RNP-donor plasmid mix was used to electroporate one million cells using the 4D-Nucleofector X Unit (Lonza) following the standard instructions and the inbuilt program (EN-130) for MCF7 cells. Post electroporation, cells were mixed with growth media and cultured following the standard procedure. Electroporated cells were FACS-sorted for mVenus-positive signal, and single-cell clones were grown in standard culture media. Single-cell clonal lines were genotyped to identify a clone correctly recapitulating the endogenous Hes1 expression. A total of 22 lines were genotyped. A clonal line (Clone 19/Cl19) with a hemizygous mVenus-Hes1 allele, passing the genotyping characterisation was chosen for all further experimentation (FigS1). For generating mScarlet-Hes1 CRISPR lines, mVenus in the donor plasmid described above was replaced with mScarlet cDNA followed by sequence verification of the new donor plasmid. The new donor plasmid was used with the same guide RNAs, following similar guidelines to generate mScarlet-Hes1 CRISPR lines (FigS4). Details of all the primers used for plasmid verification and genotyping are available on request.

**Immunostaining**

Hes1 immnostaining on MCF7 cells was performed following method as previously published (2). Hes1 primary antibody (E5 clone, sc-166410, Santa Cruz Biotech.) was used at a dilution of 1:50 in blocking solution, following incubation at room temperature for 2h. Goat anti-mouse Alexaflour (568)-labelled secondary antibody (LifeTechnologies) was used at a dilution of 1:500, following incubation for 1h. Stained cells were imaged using Olympus FV1000 confocal microscope.

**Single-molecule fluorescent in situ hybridization (smFISH) and immunofluorescence**

smFISH against Hes1 was performed as described (3) using a set of 32x20 nt probes against Human *Hes1* exonic sequence, labelled with Quasar 570 (Biosearch Technologies, US) at their 5' end. Probe details are available on request. Immunofluorescence was performed post hybridization, incubating with mouse anti-PCNA (M0879, Dako) at room temperature for 1h, and with goat anti-mouse Alexa Fluor 488 (A11001, ThermoFisher Scientific, UK) at room temperature for 30 mins.

**smFISH imaging and analysis**

Z-stack images were acquired at 0.2 μm z-increments by Delta Vision widefield microscopy (Olympus IX83 inverted microscope body, Retiga R6 2688 x 2200 pixel camera (Q-Imaging), Olympus IX3-ZDC2 laser based autofocus, 60X (1.42NA) objective, with Metamorph v7.10 (Molecular Devices) software. Identical microscopy settings were used across all images. Cellpose (4) was used to segment cell boundaries using background auto-fluorescence in the Quasar 570 channel. Segmentation masks and image stacks were combined, and cells and nuclei defined using the 'cells' module in Imaris v9.5.1 (Bitplane). Hes1 mRNA spots per cell were counted using the 'analyze spots' function in Imaris, with an estimated XY diameter of 0.5 μm. PCNA positive cells with a mean of the PCNA spot sum intensities of >80k (a.u.) were defined as S phase cells, while the ones with <80k sum intensities were defined as late G1/early S phase cells. Imaged cells showed a range of nuclear areas with linear increase in the nuclear size. Along this linear nuclear size range, lowest 20% of the cells within the smallest size range (PCNA-) were defined as G1 phase cells and 5% of the cells within the largest nuclear size range (PCNA -) were defined as G2 phase cells. M phase cells were identified on the basis of their distinct, spherical morphology.

**Protein half-life estimation**

For estimating protein half-life of exogenous HA-Hes1, MCF7 cells transfected with pCS2-HA-Hes1 plasmid were treated with 100uM Cycloheximide (CHX) for 30, 60, 90, 120, 150, 180, 210, 240, 270 and 300 mins. Cell lysates from treated cells were used for western blot analysis following standard procedures, using HA-HRP (1:1000, Roche) and tubulin antibodies (as loading control, 1:5000, Sigma).

To estimate protein half-life of endogenous mVenus-Hes1 in Cl19 CRISPR line, cells treated with CHX were live-imaged every 15 mins to capture mVenus-Hes1 mean intensities. Images were processed and analysed using IMARIS software (see below). Decreasing mVenus-Hes1 intensity values over time were used to estimate the slope and the corresponding mVenus-Hes1 half-life using Microsoft Excel.

**Incucyte growth curve analyses**

To compare the growth properties of parental MCF7 *vs* the Cl19 lines, cells were seeded on the 12 well dishes at similar cell densities in triplicates. Cells were imaged every 45 mins using the 4X lens, for around three days. Following the instructions in the Incucyte software, collected images were processed to lay a confluence mask over the growing cells, which represented cell densities over time. Percentage confluence values over time were used to estimate the growth rates of both parental MCF7 and Cl19 cells.

**Fluorescence-activated cell sorting (FACS) to obtain subpopulations of cells from a heterogeneous mixture**

To obtain $CD44^{High}CD24^{Low}$ breast cancer stem cells (bCSCs) and bulk (non-stem) subpopulations, cells were trypsinised, resuspended in DMEM/10% FBS and counted using a Neubauer chamber. Ten million cells per FACS sort were centrifuged (500g, 3min), resuspended in blocking solution (1X PBS with 2% FBS, 1000µL) and incubated on ice (30 min). Post blocking, cells were centrifuged again (500g, 3min). Blocking solution was replaced with 100µL blocking solution containing appropriate antibodies (allophycocyanin-conjugated CD44/CD44-APC, Beckton Dickinson and phycoerythrin-conjugated CD24/CD24-PE, Beckman Coulter) at 1:40 and 1:80 dilutions, respectively. Cells were incubated on ice (30-60min). Post-incubation, cells were washed twice with PBS before resuspending in 1X Hank's Balanced Salt Solution (HBSS) at a concentration of five million cells/ml containing 10µg/ml DAPI. Stained cells were FACS sorted to obtain purified $CD44^{High}CD24^{Low}$ bCSCs and bulk cells from the MCF7 lines using a BD FACSAria Fusion cell sorter (BD Biosciences) as described before (5). To obtain $ALDH^{High}$ bCSCs, cells were stained using AldeRed ALDH Detection Assay kit (Sigma) or ALDHFLOUR kit (STEMCELL Technologies), following the instructions from the manufacturer. Stained cells were FACS sorted using FACSaria cell sorter into $ALDH^{High}$ bCSCs and $ALDH^{Low}$ bulk cells (6).

**Blocking cell-cycle using $CoCl_2$** To block cell-cyle in MCF7 reporter cells, we treated them with $CoCl_2$, a known hypoxia memetic (7). $CoCl_2$ (Merck, 255599) was suspended in sterile water. Cells were treated with 300uM $CoCl_2$ for 2-3 days to block their cell-cycle completely, before we started imaging the reporter lines as described in the section below. Control cells were treated with equal volumes of sterile water.

**Data collection using confocal microscopy and cell tracking using IMARIS**

MCF7 lines expressing fluorescent reporters were live-imaged every 15 mins (unless stated otherwise) for 60-120hr using a Nikon A1 confocal microscope. The resulting time-lapse, Z-stacked images were spot-tracked manually using image analysis software IMARIS to obtain 'mean' intensity profiles for Hes1 expression, which represented changes in Hes1 'concentration' during that time, and also to identify precise locations of cell mitoses and various phases of cell-cycle.

The Hes1 mean intensity levels (in arbitrary units/a.u.) obtained from IMARIS were plotted against time for single cells to obtain single-cell dynamic Hes1 traces over time (time tracks, Fig1D and FigS4C, 7A) for the entire duration of the movie. The entire Hes1 intensity data could be plotted in the form of cell lineage trees (FigS7B). These lineage trees not only showed the changes in the

Hes1 intensity over the entire duration of the movie but also exhibited these changes across cell generations, consisting of mother and daughter cells. In majority of the cases, we extracted the mean Hes1 intensity changes over time between consecutive mitoses, representing full cell-cycles for two cell generations (G1 and G2, FigS7B). These data formed the basis for our analyses of our experiments.

**Data analyses methods and custom generated codes**

All data analyses were performed on MATLAB R2018a unless stated otherwise. All the custom codes/algorithms used for data analyses have been deposited in the following GitHub repository: https://github.com/AndyRowntree/Interaction-of-Hes1-oscillations-with-cell-cycle.

**a) Generation of temporal Hes1 expression profiles as time tracks and heat maps**

To enable the comparison of large numbers of dynamic time traces in one graph, we decided to display dynamic traces from individual cells as normalised heatmaps. First, due to high heterogeneity in the Hes1 mean intensity levels, intensity values over time from each cell were normalized using z-scores (using the formula $y = (x - \mu)/\sigma$, where y and x represent the normalised and raw intensity levels, respectively, at each time point. $\mu$ and $\sigma$ are the mean and standard deviation, respectively, of the intensity levels across the whole trace) to provide relative traces (as shown in FigsS7). We then used these normalised Hes1 intensity values to generate heatmaps using the '***imagesc'*** function, where each x-axis point contains one cell's dynamic Hes1 trace from the first mitosis to the second mitosis (representing a complete cell-cycle) with time running vertically from top to bottom.

**b) Interpolation**

In order to classify the heat maps of varying lengths using clustering algorithms, we needed to take each time series and display it as a pseudo-time series of a constant chosen length. We 'stretched' or 'squashed' the data to a pseudo-time scale using the default interpolation option (linear interpolation) using the '***interp1***' function in MATLAB. Due to the requirements of the GMM clustering algorithm (see below), which requires more columns than rows in order to perform clustering, the number of time points in the pseudo-time scale was set to the number of traces minus one. However, when displaying the pseudo-time traces as heatmaps (as shown in Fig6), this number was increased to the length of the longest trace so to encompass more detail into the heatmaps. To ensure that no clusters were produced as artefacts due to this constraint of pseudo-time points, we performed a synthetic control (see below).

**c) Gaussian Mixture Model (GMM) clustering**

Clustering is a standard data analysis technique, classifying data into distinct groups, each containing a specific trait or characteristic. We performed a Gaussian mixture model (GMM) clustering to classify Hes1 heat maps into discernible, individual groups (8). This clustering method fits unlabeled data into a predetermined number (k) of d-dimensional Gaussian components (in our case, d is the number of pseudo-time points). Albeit somewhat similar to k-means clustering, GMM clustering allows for more flexibility as both the mean and variance/covariance of components are considered instead of merely the means. GMMs also allow for distributions, which may be elliptical and not strictly spherical like k-means, allowing for more freedom. We utilised the functions '***fitgmdist***' and '***cluster***' to perform our clustering, it fits Gaussian components to the data using the iterative Expectation-Maximisation (EM) algorithm. Random initial values for component means, covariance matrices, and mixing proportions are chosen, and these parameters are tuned through each iteration until a desired convergence is reached. We use the Silhouette method (9) to determine the optimal number of clusters (k=3) . This method provides a value (the silhouette coefficient) which can be used as a measure of how similar each trace is to its assigned cluster and to other clusters. A score (per k) can then be obtained through averaging of these values: the higher the score, the more suitable it is for the data to be classified into k clusters. The method was also used to suggest the most suitable constraints to place upon covariance matrices of the Gaussian components namely 'shared', which means that the shape of distributions for all components in the mixture model are identical and 'diagonal', meaning that Gaussians distributions were only allowed to stretch or squeeze in orthogonal directions during the EM algorithm iterations (FigS8A-B).

**d) Synthetic Data as a control for GMM clustering**

When interpolating data using a pseudo-timescale, the shape of the Hes1 dynamic expression may be distorted and features in cells with longer cell-cycles might be missed. To ensure that our clustering techniques would perform correctly on our data set and clusters aren't created simply as a result of stretching/squashing, we constructed synthetic data as a control. For this we generated five underlying dynamic shapes: three from polynomial curves fitted to our actual data, and two antiphase sine curves (FigS8C). These antiphase sine curves would generate two sets of traces that were directly opposed and therefore should never cluster together. Each shape was generated over short, medium and long length-scales in order to produce 15 base traces (FigS8D). We then duplicated each base trace ten times and added a vector X of random values, where $X \sim \mathcal{N}\left(0, \frac{1}{2}\right)$ to represent biological noise, which was representative of our actual experimental data, resulting in 150 (n = 152 in our actual experimental data) distinct, synthetic control traces. We then interpolated these traces to provide data of the same length, followed by randomly mixing the (FigS8E) and applied the same clustering techniques as with our previous analysis. Results showed that all 5 dynamic shapes were clustered together (FigS8G) using GMM clustering at a cluster number and covariance matrix type as suggested by silhouette analysis (FigS8F). These results confirmed that the clustering methods used in our experimental analysis were not skewed as a result of data interpolation.

**e) Automated Hes1 expression dip-detection pipeline**

To detect a dip in the normalised mVenus-Hes1 expression during the cell-cycle in an automated fashion (Fig4 E,F and 5B, C), we developed a computational pipeline, into which our data consisting of the Hes1 mean expression time traces were fed. For each Hes1 trace, Savitzgy-Golay (SG) filter was applied using the function '***sgolayfilt***'. Parameters for the SG filter were set as a polynomial order of 3 and a moving frame length of a quarter of the given trace length. The resultant smoothened traces were truncated by three time points at either end to remove artefacts caused by the low polynomial order. All turning points of smoothened, truncated traces were found by finding the points at which the product of the difference between these trace values at three consecutive time points is negative. The dip in expression was then determined as being the point at which the minimum of all such turning points lied. To avoid earlier detection of a dip in any cells starting with low Hes1 levels, dips detected in the first ¼ part of any trace were neglected. The same smoothened traces were utilised when finding the Hes1 peak in G1 phase (Fig7A): this was accomplished by identifying the maximum value of our traces which lay earlier in the cell-cycle than our previously-given dip.

**f) Periodicity Analyses of Hes1 time traces**

Visual inspection of our mVenus-Hes1 time traces suggested that Hes1 expression was oscillatory with a single 'full' wave encompassing more than one cell-cycle/generation. Hence, we found it difficult to reliably extract periodicity data for the Hes1 wave from time traces covering only a single cell-cycle. To confidently extract the periodicity values of a Hes1 wave, using more than one analysis pipelines (see below), we concatenated Hes1 time traces for two generations (combining Hes1 mean intensity traces from two generations, mothers and their daughters) (n=164). Our analyses across generations also showed that there was no loss of Hes1 protein during mitoses connecting these two cell-cycles/generations. The mean Hes1 levels at the end of the first cell-cycle and the beginning of the second cell-cycle were nearly identical (Fig2E); the overall Hes1 protein levels during mitosis were also matching to the Hes1 levels in mother and daughter cells (data not shown). These observations confirmed the continuation of a single Hes1 wave across generations, without any loss of Hes1 protein. These periodicity analyses were performed using a variety of methods for corroborative reasons and are described below.

**1 - Continuous-wavelet based time-frequency analysis of Hes1 periodicity using pyBOAT**

To correctly identify time-dependent properties of oscillations within our Hes1 time-series data, we used pyBOAT (10). PyBoat is a computational tool to identify and characterise oscillations in time series data. Its code, detailed installation and user guidelines are available on the GitHub (https://github.com/tensionhead/pyBOAT). It includes built-in functions for Fourier transforms and continuous wavelet transforms. Wavelet transforms can be particularly useful to identify any time-dependent features of non-stationary oscillatory signals. Before Fourier or Wavelet transforms are applied, pyBoat detrends each time series. This detrending relies on a windowed sinc filter (11), which is designed to remove all Fourier components with periods longer than a fixed threshold, $T_c$, from the signal. In all our analysis, we chose $T_c$ equal to the maximum length of the considered time series. Typically, our time series covered a range of 3000-4000 mins, covering two cell-cycles/generation for the MCF7 cell subpopulations imaged. When applying wavelet analysis, we consider periods between 1h and the maximum length of each track.

**2 - Lomb-Scargle periodogram (LSP)**

The Lomb–Scargle periodogram (12) is a well-known algorithm, falling under the Fourier transform and least-squares methods for detecting and characterising the periodicity from time-series data. It allows efficient computation of a Fourier-like power spectrum estimation, resulting in an intuitive means of determining the period of oscillation. We utilised the '***plomb***' function within MATLAB to estimate periodicity via this method for comparison with our Ubc:mVenus-Hes1 control (Fig 10). It returns the Lomb-Scargle power spectral density (PSD) estimate of a signal that is sampled at the instants specified. We input linearly detrended traces and received a normalised (MATLAB allows for a 'normalize' specification with the '***plomb***' function which outputs a power value between 0-100 for ease of comparison between experiments) periodogram as an output. One particularly high normalised power spike from these spectra imply a strong oscillatory signal. In contrast, readouts consisting of multiple spikes which are relatively lower in power implies the lack of oscillation in our traces (exemplified in Fig 10 B).

**3 - Autocorrelation**

We used the '***xcorr***' function to determine a periodicity estimate via autocorrelation. This method finds the cross-correlation (13) (value of a given trace against itself as it is shifted along in time. The autocorrelation sequence of a periodic signal has the same cyclic characteristics as the signal itself. Thus, autocorrelation can help verify the presence of cycles and determine their durations. When more than one period was recorded per trace, we took the mean of the recorded period estimates that satisfied a pre-set threshold determined via bootstrapping; this ensured that oscillations weren't a result of random noise.

**4 - Ultradian periodicity analyses**

We ran our time series data through the Gaussian processes based pipeline developed in our lab to detect ultradian oscillations, following the guidelines described in the paper (14). In brief, the method combines mechanistic stochastic modelling with methods of non-parametric regression with Gaussian processes to distinguish ultradian oscillations from random fluctuations. MatLab script for the same is available through the GitHub repository.

**g) Phase diagrams**

We define phase of an oscillation as an angle between 0 and $2\pi$ which corresponds directly to a cosine wave (Fig 8 A bottom panel) between the same values (i.e. phase at $0=2\pi$ is a peak and phase at $\pi$ is a trough in the wave). To extract a phase progression readout, we applied the Hilbert transform onto smoothened normalised (z-scored) mVenus-Hes1 pseudo-time traces from mitosis to mitosis (Fig 8 A top panel). We used MATLAB functions '***hilbert***' and '***angle***' to output the given phase at all points in pseudo-time for each trace (Fig 8 A middle panel). Smoothening was performed using the function "***smoothdata***" and with the specifications of "***gaussian***" smoothing and a moving window of 20 pseudo-time points (whole traces were stretched to 144 time points). Polar scatter plots were plotted using the function '***polarscatter***' (Fig8C and D). They represented the phase of each Hes1 wave at the beginning of mitoses, showing phase outputs for each trace at the start of a new cell-cycles.

**Correlation plots**

We used the Spearman correlation coefficient plots to measure the strength and direction of a linear relation between two variables (such as Hes1 mean intensity and cell-cycle length). We used the following textual descriptors associated with R values to describe the relationship between our variables; R = 0 means no linear relationship, +0.30 a weak positive linear relationship, +0.50 a moderate positive relationship. +0.70 a strong positive relationship and 1 a perfect positive relationship. Also note that for a given R value, its square value denotes the proportion of the variance that can be attributed to the correlation; for example, for a correlation between cell-cycle length and mean Hes1 intensity, an R value of 0.33 and its squared ($R^2$) value of 0.11 would mean that only 11% of the variance in the cell-cycle length can be attributed to its correlation with the Hes1 levels, clearly showing why an R value of 0.33 represents a weak positive correlation.

**Statistical analysis**

Data were analysed using multiple hypothesis testing methods, as indicated in the figure legends using Prism 8 (GraphPad, USA). Statistical significance is reported for p-values <0.05 (*), <0.01(**) and <0.001 (***), calculated using unpaired T tests. Errors bars are reported as standard deviation of the mean (mean+/-SD) calculated from pooled data unless otherwise stated. We also used

Bartlett's test (through MATLAB) to check if the variances were equal across the groups of data. Significant Bartlett statistic allowed rejection of the null hypothesis of equal variances across groups.

**Legends for supplementary figures**

**FigS1. Genotyping of the selected CRISPR clones by PCR, followed by DNA sequencing confirmed the correct insertion of mVenus cassette at the Hes1 N-terminal in Cl19 cells.** (A) Schematic of Hes1 genomic locus, pre- and post-tagging with mVenus. Forward and reverse primers (FP and RP) used for PCR-based genotyping have also been designated. (B) PCR results using FP and RP performed using genomic DNA isolated from different CRISPR clonal lines and the parental MCF7 cells as a control. Take note of a single, smaller band (wild type allele without any insertion) with parental MCF7 cells. Clonal lines 7 and 19 showed two PCR bands, with the upper/larger one suggesting amplification from the Hes1 allele after mVenus insertion; clone 11 also showed a diffused upper band. Both PCR bands from clones 7, 11 and 19 were cloned in TOPO vectors and sequence verified. (C) Sequencing results for the smaller PCR band (allele without insertion) from clone 19. Sequencing data showed substitution and insertion mutations near the PAM sites, resulting in the introduction of a premature stop codon, causing knocking-out of this Hes1 allele. (D) Sequencing results for the larger PCR band (allele with insertion) from clone 19. Sequencing showed the insertion of 3Flag-mVenus-Linker-HA cassette immediately after the start codon of Hes1 coding region. Sequencing data (C and D) confirmed that clone 19 is hemizygous for Hes1, with one allele knocked-out and the other one with the correct insertion of the mVenus cassette.

**FigS2. Incucyte based growth comparisons showed that the mVenus-Hes1 CRISPR clonal line (Clone 19/Cl19) and its parental MCF7 line had similar growth properties.** (A) Selected images at different time points from the cells (parental MCF7, upper panel and Cl19, lower panel) growing in the incucyte incubator. Areas with cells growing were masked using the incucyte software to measure the percentage confluence, which, when plotted against time measured the growth rates for both cell lines. The growth rates (dark blue trend lines on the graph) for parental MCF7 and Cl19 cells were identical (B), suggesting that losing a Hes1 allele did not affect the growth properties of Cl19 cells.

**FigS3. Protein half-life comparisons between mVenus-Hes1 from the Cl19 cells and HA-Hes1 over-expressed in the parental MCF7 suggested that tagging Hes1 with mVenus did not affect its half-life.** (A) Time course of live-imaging of Cl19 cells while they were treated with either ethanol/EtOH (control, upper panel) or cycloheximide/CHX (lower panel). (B) Hes1 mean intensities over the time course of EtOH/CHX treatment for single cells, tracked using IMARIS software. The slops of these intensity plots were used to estimate protein half-lives in Excel. (C) Half-lives of more than 100 cells, which were imaged while being treated with CHX, with a mean half-life of around 4hr. (D) Western blot analysis performed on lysates from MCF7 cells over-expressing HA-Hes1 and treated with cycloheximide/CHX (insight panel). The main panel shows Hes1 protein-life, estimated from the measured band intensities from the western blot, to be 4.1h. These two independent approaches to measure protein half-lives suggested that tagging Hes1 with mVenus did not affect its half-life.

**FigS4. (A-C) Additional clones where Hes1 was endogenously tagged with mScarlet through CRISPR/Cas9 reaction showed similar dynamics to the mVenus-Hes1 C19 line. (D) Existence of the endogenous Hes1 protein dynamics was strengthened by the observation that the numbers of Hes1 mRNA were not constant through the cell-cycle as observed by the smFISH analysis on parental MCF7 and Cl19 cells.** (A) Schematic of Hes1 genomic locus before and after CRISPR/Cas9 based insertion of mScarlet cDNA prior to exon 1. Primers used for genotyping have been designated as half arrows. (B) Genotyping summary of two mScarlet-Hes1 CRISPR/Cas9 clones, used to corroborate our findings that Hes1 shows temporal dynamics, as deduced from our analyses using mVenus-Hes1 time traces from another CRISPR line (Cl19, expressing mVenus-Hes1) (Figs1D, 3, 4 and 5). One of these clones is hemizygous for Hes1 (M18) and the other one (Cl12) is heterozygous. (C) Example time traces for mScarlet-Hes1 from M18 and Cl12 cells, collected by the single-cell live-imaging of these lines. (D) Example images of Hes1 smFISH on parental MCF7 cells, separated for different cell-cycle phases. Graphs underneath shows that Hes1 mature mRNA numbers through the cell-cycle are not constant; G2 phase cells have higher Hes1 mRNA than G1 phase cells for both MCF7 and Cl19 cells.

**FigS5. Power spectrum analyses of Hes1 wave using LSP platform** (A) Example of endogenous mVenus-Hes1 (left panel) and exogenous Ubc:NuVenus (as a non-oscillatory control, middle panel) traces were linearly detrended and run through the Lomb-Scargle periodogram/LSP pipeline, which gives normalised (scaled between 0-100) power spectrum for the input traces (B). The dominant frequency and the corresponding period values were extracted from these power spectra. Notice a dominant, high peak associated with the mVenus Hes1 trace, compared to multiple, lower peaks associated with the control trace. (C) Normalised power values for the dominant periods for experimental mixed cells were significantly higher in comparison to the Ubc:NuVenus control cells.

**FigS6. Analyses of ultradian periodicities in mVenus-Hes1 traces show that all cells (irrespective of their cluster types or cellular subtypes) show ultradian periodicity of around 6h with low mean fold-change values, embedded in the larger, circadian-like periodicity of Hes1. Neither the periodicity nor the fold change showed any correlation with the cell-cycle length.** (A) Top panel shows an example of single cell mVenus:Hes1 trace (thin red line) with trend line superimposed (thick red line). Bottom panel shows the same raw trace with the trend removed (thin blue line) and a new oscillatory trend superimposed on top (thick blue line). (B) Boxplots of ultradian periodicities from passing oscillators (as determined with a Gaussian Process algorithm, see m&m), separated as per cellular subtypes (left) and cluster classification (right). (C) Distribution of mean ultradian periodicities of all cells (left), showing no correlation with the cell-cycle length (right). (D) Distribution of mean fold change values (amplitude) for ultradian periodicities of all of all cells (left), showing no correlation with the cell-cycle length (right).

**FigS7. Representing Hes1 intensities as heat-maps and cell trajectories as cell-lineage trees.** To compare multiple, single-cell Hes1 time traces against one another, we plotted Hes1 mean intensities over time as heat maps. (A) Example showing how an mVenus-Hes1 single-cell time-trace is converted into a heat map after normalising the Hes1 intensity. Refer to materials and methods (m&m) for details about how intensity values are normalised. (B) MCF7 cells are highly amenable to single-cell live-imaging approaches. Cells can be live-imaged across generations and single cells can be tracked to generate lineage trees. B shows how a lineage tree is generated by imaging a cell across generations, through mitoses/branching points, following its daughters and grand-daughters. Information about Hes1 levels, dynamics and cell-cycle kinetics were obtained for generations with a full cell-cycle, as in this lineage tree for generation 1.

**FigS8. Silhouette analysis, followed by the GMM clustering analysis on the synthetic data justified the number of GMM cluster choice.** A) Line graph of means of multiple, repeated Silhouette scores for GMM clustering at four covariance matrix types (shared and full, shared and diagonal, unshared and full, unshared and diagonal) with number of clusters ranging from 2 to 8. The image shows that the optimal GMM clustering for our data (i.e. the highest point on the graph) was a 3-component mixture with shared (identical covariances) and diagonal (only orthogonally flexible Gaussian components) covariance matrices. (B) Line graph of the orange line in (A) showing again the mean but with +/- one standard deviation error shading of multiple repeats of such clustering. This shows that the optimal component number and covariance matrix type were consistent amongst multiple repeats of the Silhouette score analysis. (C) Five underlying dynamic shapes: 1, 2 and 3 based on dynamics of actual data and 4, 5 sine and negative sine waves respectively. (D) Heatmap showing 10 noisy (representing biological noise) duplicates of each dynamic shape trace at three different time length scales (representing cell-cycle length heterogeneity). (E) Heatmap of stretched (using identical interpolation as with experimental data) and randomly ordered synthetic traces from (D). (F) Line graph of means of multiple, repeated Silhouette scores for GMM clustering at four covariance matrix types (shared and full, shared and diagonal, unshared and full, unshared and diagonal) with number of clusters ranging from 2 to 8. The image shows that the optimal GMM clustering for our synthetic data was a 5-component mixture with shared diagonal covariance matrices. (G) Heatmap showing GMM clustering using optimal cluster number and covariance matrix type as suggested by Silhouette analysis. This image shows all five original dynamic shapes being clustered together, thus eliminating the possibility of stretching data producing artificial clusters. These control data together clearly show that clustering is not an artefact of data stretching and the numbers of clusters suggested by Silhouette analysis are correct.

**FigS9. GMM clustering performed on time-stretched data followed by showing the clustered data in an unstretched form,** confirmed that the normalised Hes1 mean expression patterns falling into three distinct clusters was not an artefact of stretching the Hes1 traces.

**FigS10. Mean Hes1 traces and mean cell-cycles mapped across two cell-cycle generations illustrate how the dynamics of Hes1 and the cell-cycle lengths transition when cells move across different clusters.** (A) Mean Hes1 intensity (in dark blue with standard deviation in lighter blue, shaded) for all cells across two-generation traces in pseudo-time of all cells falling into 9 cluster transitions with division points marked as a red dot. This illustrates the transition in dynamics across generations and elucidates the cluster switching across two generations. (B) Boxplots of all cell-cycle durations (gen 1 on left and gen 2 on right within each panel) for all 9 cluster transitions. The data shows elongation, shortening or duration-maintenance of cell cycles between mother and daughter cells depending on their cluster transitions (as viewed in A). Note that transitions into cluster 3 elongate the cell cycle, while transitions out of cluster 3 shorten it.

**FigS11. Periodicity analyses across two cell generations for possible cluster transitions suggest that cluster transitions involving cluster 3 (transition from or to cluster 3) involves a change in Hes1 periodicity across generations, and a corresponding change in the cell-cycle length.** (A) Distribution of the periodicities of mVenus-Hes1 traces as histograms show the range of periodicities shown by mVenus-Hes1 single-cell time tracks. (B) Mean periodicity over time (dark blue lines) with standard deviation (in light blue shades) for all traces for two generation in pseudo-time arranged via cluster to cluster transitions (9 transitions). The data show an increase in periodicity after a division when cells transition from cluster 1 to cluster 3 transition, and a decrease in periodicity after cell division when cells transition from cluster 3 to cluster 1. (C) Scatter plot of the mean periodicity for each trace over two generations against the cell-cycle durations of the daughter cells (in second generation) showed moderate correlation among the two, suggesting an increase in the cell-cycle length when there is a corresponding increase in the Hes1 periodicity. (D) Example mVenus:Hes1 trace of two generations stitched together in which both generations are classified as cluster 1 (left). Example mVenus:Hes1 trace of a two generations together stitched together, in which the first generation is classified as cluster 1 and the second as cluster 3 (right).

**FigS12. Over-expression of Hes1 under sustained promotes the CD44$^{High}$CD24$^{Low}$ type of cell fate.** (A) Example of FACS analysis of MCF7 cells stained with CD44-APC and CD24-PE antibodies (see details in m&m). Live gated cells (top left panel) are gated further in a way that the Q4 quarter represents around 5% of CD44$^{High}$CD24$^{Low}$ cells for control cells (parental MCF7). (B) When the same gates were applied to the transduced cells (Ubc:NuVenus or Ubc:mVenus-Hes1), the proportion of CD44$^{High}$CD24$^{Low}$ cells expressing NuVenus was similar to untransduced cells but cells expressing Ubc:mVenus-Hes1 showed significant increase in the CD44$^{High}$CD24$^{Low}$ population.

**FigS13. Exogenous Hes1 expressed under the Ubc promoter does not raise the total Hes1 level because it suppresses the endogenous Hes1 expression**. (A) Upper panels show endogenous mVenus-Hes1 expression (Cl19) and middle panel shows endogenous mScarlet-Hes1 expression (M18). Lower panel shows images of M18 cells transduced with exogenous mVenus-Hes1 under the Ubc promoter, showing how exogenous mVenus-Hes1 has taken over the endogenous expression of mScarlet-Hes1. Scale bar=50um. (B) shows the quantification of overall Hes1 levels (combination of mVenus-Hes1 and mSCarlet-Hes1) for untransduced Cl19, M18, and transduced M18 with Ubc:mVenus-Hes1 cells, showing that overall Hes1 levels in the transduced cells are comparable to the untransduced cells. (C) The overall Hes1 level in transduced cells (M18 cells expressing mScarlet-Hes1 transduced with Ubc:mVenus-Hes1) is mainly contributed by the exogenous Hes1 (Ubc:mVenus-Hes1), with almost zero contribution from the endogenous Hes1 (mScarlet-Hes1), which is suppressed.

**FigS14. Overexpression of sustained Hes1 suppresses the dynamics of p21 expression.** (A) Images extracted from single-cell live-imaging of p21-mVenus CRISPR MCF-7 cells, transfected with either (A) control plasmid (Ubc:mScarlet) or (B) experimental plasmid (Ubc:mScarlet-Hes1). mScarlet signal is shown in red and p21 signal is shown in green. Right panel; single-cell tracks show that in cells transfected with the control plasmid, p21 shows dynamic expression, with a prominent peak in late G1 phase. In cells expressing mScarlet-Hes1 under the sustained (Ubc) promoter, dynamics in p21 expression were lost, with cells showing sustained p21 signal and no cell division. (C) For cells with full cell-cycle, when we plotted p21 intensity against the cell-cycle length, no correlation was observed, suggesting that the levels of p21 do not affect the length of the cell-cycle. While around 75% of cells with dynamic p21 showed cell divisions, by contrast, around 75% of cells with non-dynamic p21 showed no division (D). C and D together suggest that the dynamics of p21 is important for cell-cycle progression.

**Legends for supplementary movies**

**Supplementary movie 1** shows time-lapse images (15fps) of the Clone 19 reporter line wherein the clonal cells express endogenously tagged Hes1 with mVenus (green). These cells were imaged every 15 mins for a duration covering at least one complete cell-cycle (mitosis to mitosis) of FACS-sorted cellular subpopulations from MCF7 cells, including $CD44^{High}CD24^{Low}$ breast cancer stem cells (bCSCs), $ALDH^{High}$ bCSCs and non-stem bulk cells. This line reported the dynamics of the endogenous mVenus-Hes1.

Clone 19 cells were further transduced with the lentiviral cell-cycle reporter (mCherry-PCNA, red). This new line, shown as **supplementary movie 2** (**supplementary movie 2a** showing mVenus-Hes1 signal in green and **supplementary movie 2b** showing mCherry-PCNA signal in Red for the same set of cells) formed the basis of our analyses connecting dynamic protein expression of mVenus-Hes1 with different phases of the cell-cycle.

### Abstract in picture

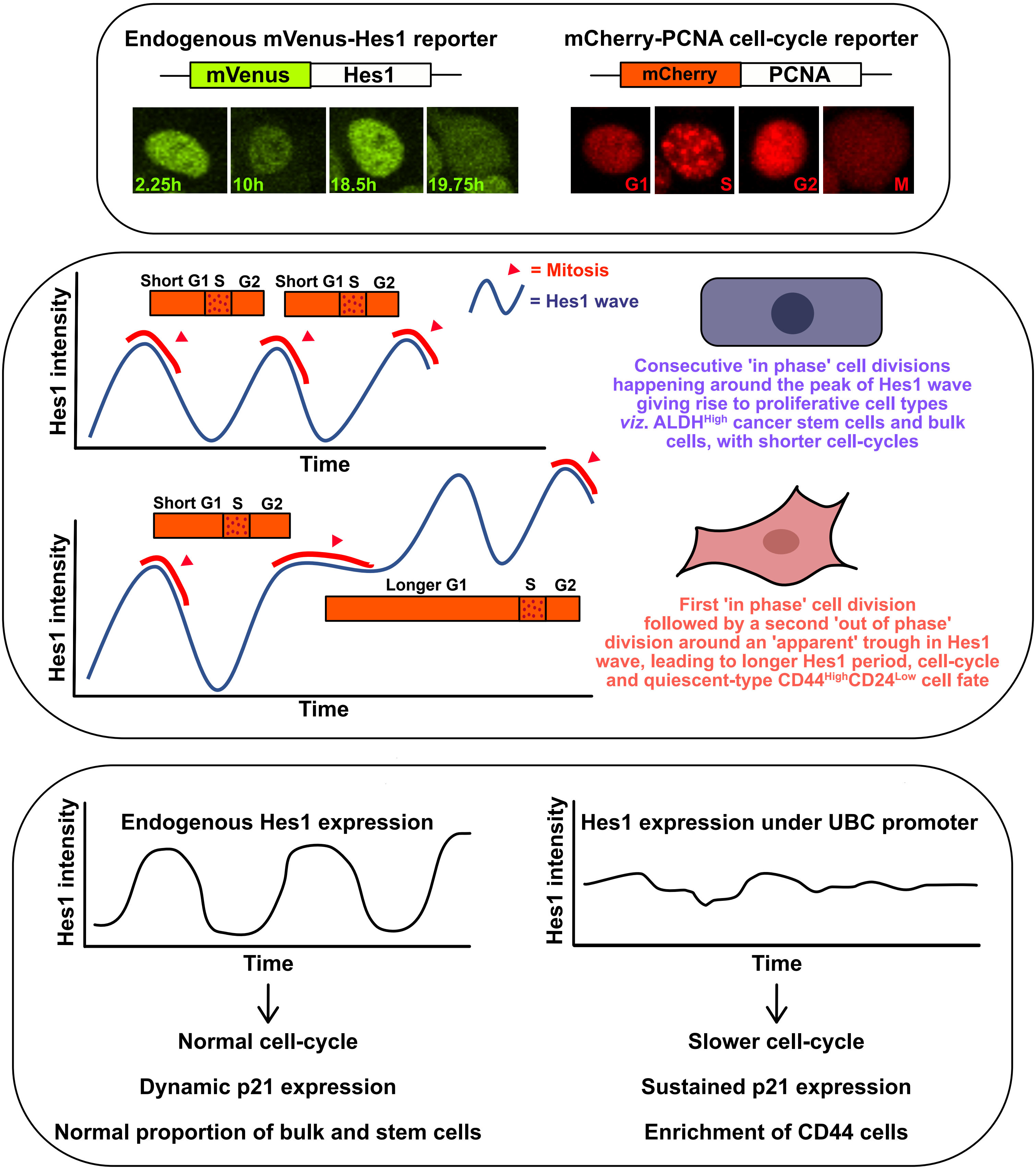


Figure S0

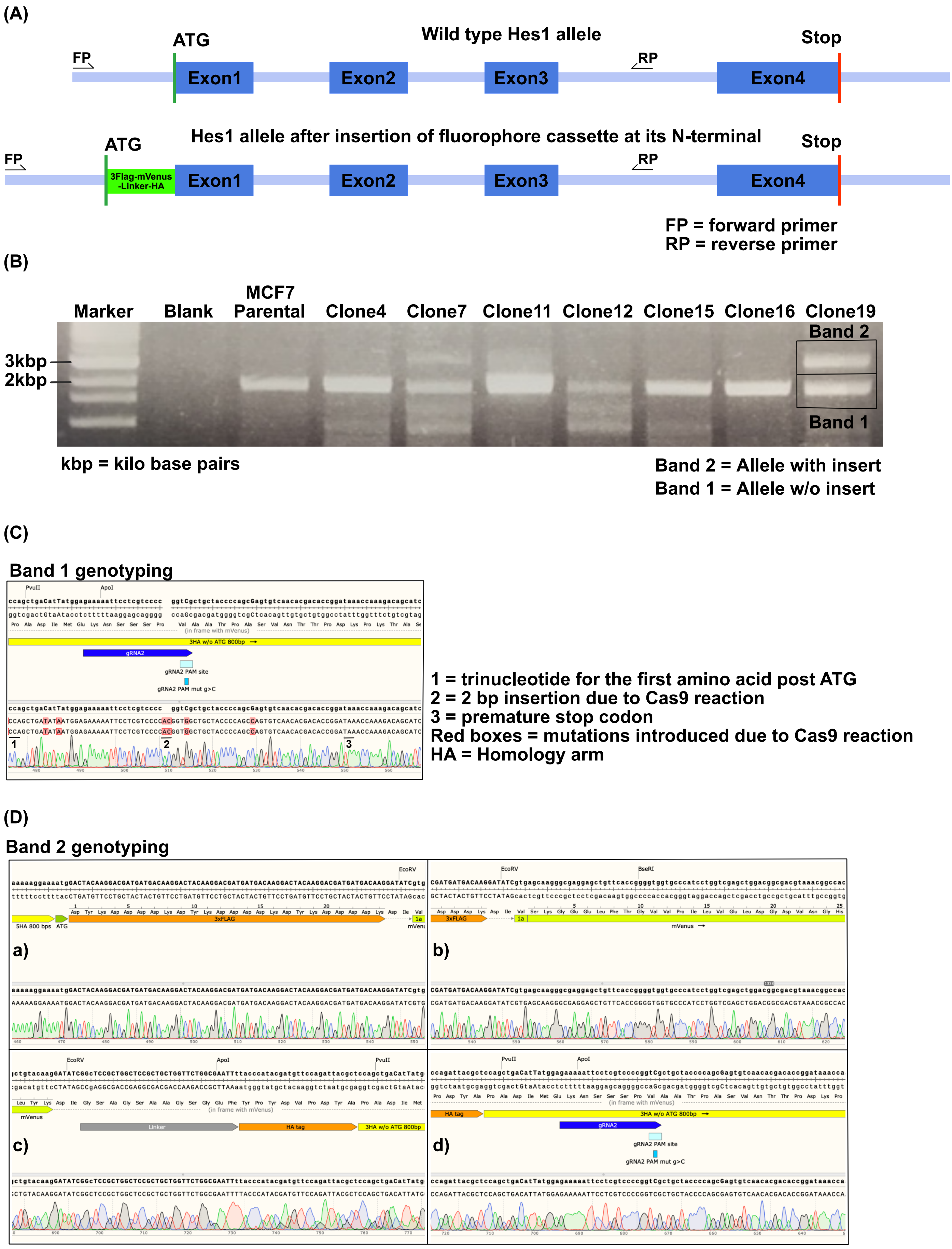
(A)
Wild type Hes1 allele
FP
ATG
Exon1
Exon2
Exon3
RP
Stop
Exon4
Hes1 allele after insertion of fluorophore cassette at its N-terminal
3Flag-mVenus -Linker-HA
FP = forward primer
RP = reverse primer
(B)
Marker
Blank
MCF7 Parental
Clone4
Clone7
Clone11
Clone12
Clone15
Clone16
Clone19
Band 2
Band 1
3kbp
2kbp
kbp = kilo base pairs
Band 2 = Allele with insert
Band 1 = Allele w/o insert
(C)
Band 1 genotyping
1 = trinucleotide for the first amino acid post ATG
2 = 2 bp insertion due to Cas9 reaction
3 = premature stop codon
Red boxes = mutations introduced due to Cas9 reaction
HA = Homology arm
(D)
Band 2 genotyping
a)
b)
c)
d)

Figure S1

(A)

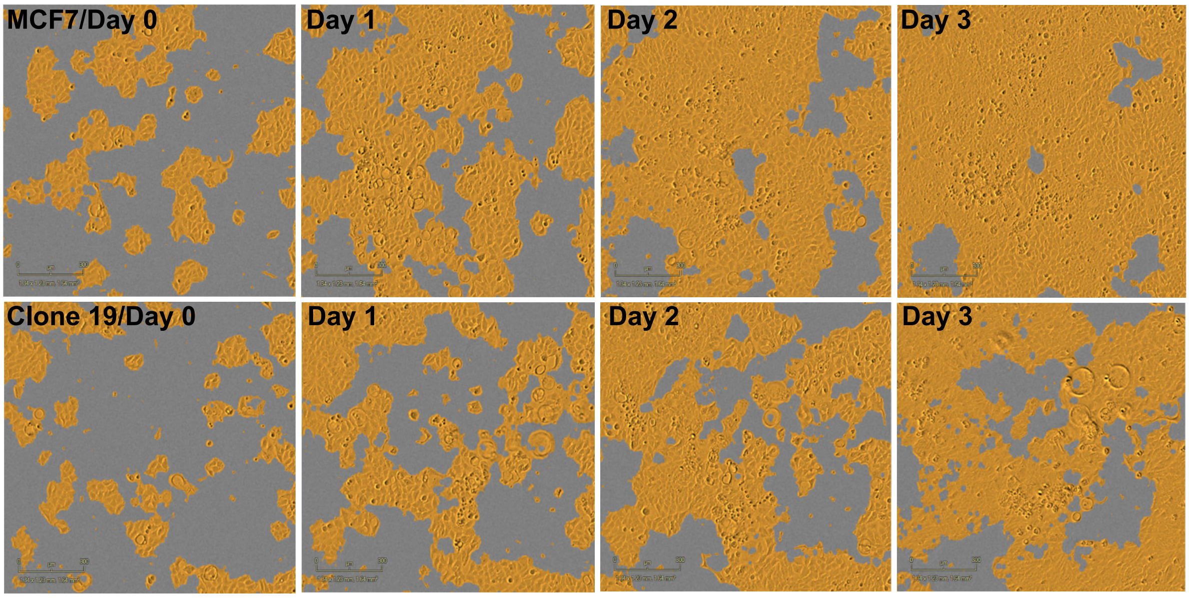
MCF7/Day 0
Day 1
Day 2
Day 3
Clone 19/Day 0
Day 1
Day 2
Day 3

(B)

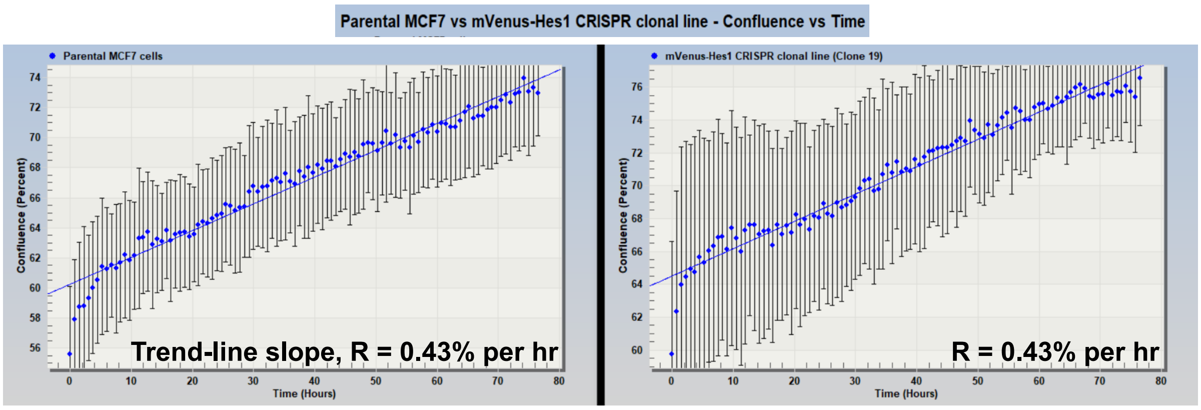
Parental MCF7 vs mVenus-Hes1 CRISPR clonal line - Confluence vs Time
Parental MCF7 cells
Confluence (Percent)
Time (Hours)
Trend-line slope, R = 0.43% per hr
mVenus-Hes1 CRISPR clonal line (Clone 19)
Confluence (Percent)
Time (Hours)
R = 0.43% per hr

Figure S2

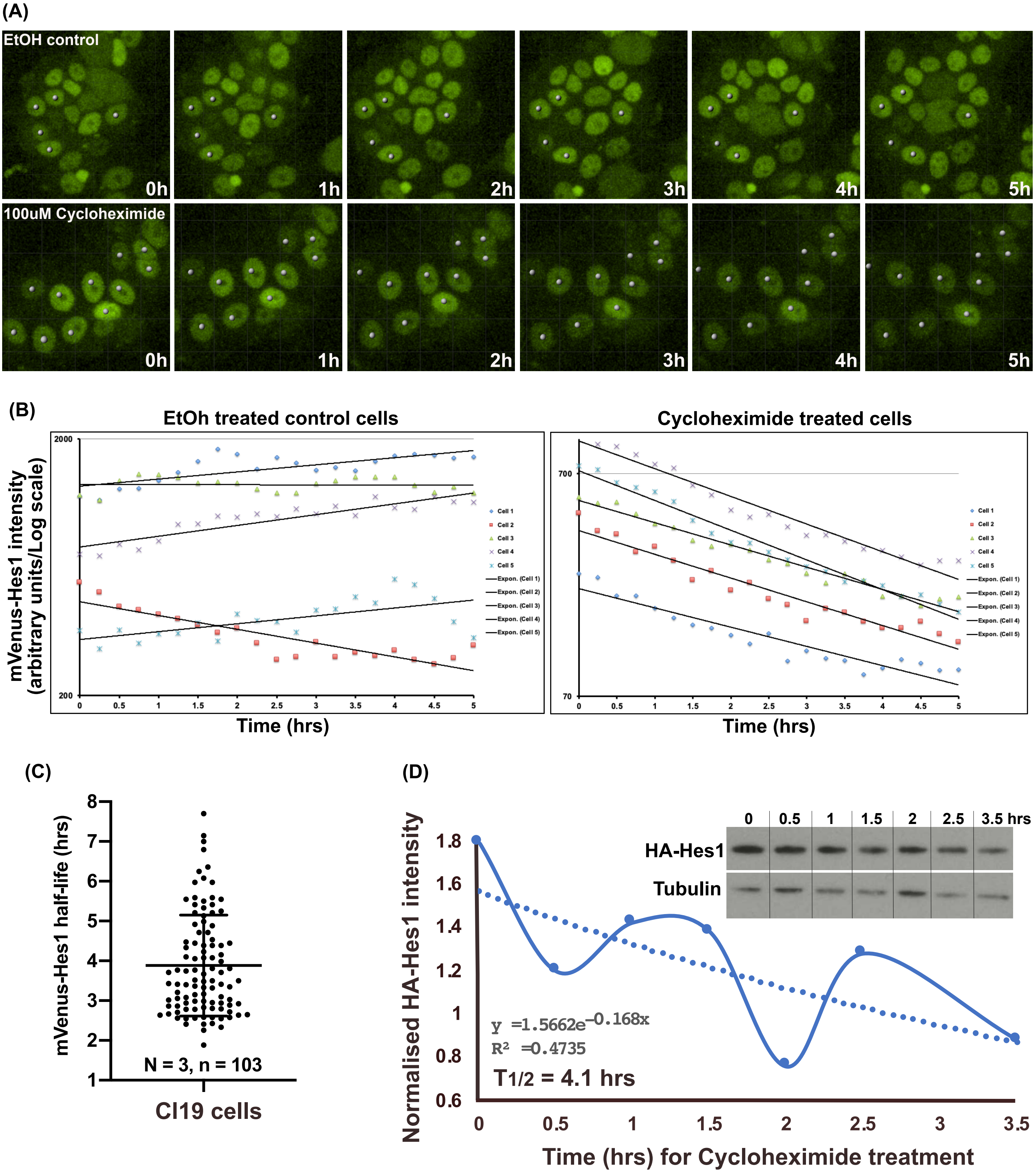

(A)
EtOH control
0h
1h
2h
3h
4h
5h
100uM Cycloheximide
(B)
EtOh treated control cells
Cycloheximide treated cells
mVenus-Hes1 intensity (arbitrary units/Log scale)
Time (hrs)
Cell 1
Cell 2
Cell 3
Cell 4
Cell 5
Expon. (Cell 1)
Expon. (Cell 2)
Expon. (Cell 3)
Expon. (Cell 4)
Expon. (Cell 5)
(C)
mVenus-Hes1 half-life (hrs)
N = 3, n = 103
Cl19 cells
(D)
Normalised HA-Hes1 intensity
0 0.5 1 1.5 2 2.5 3.5 hrs
HA-Hes1
Tubulin
y =1.5662e−0.168x
R² =0.4735
T1/2 = 4.1 hrs
Time (hrs) for Cycloheximide treatment


Figure S3

(A)

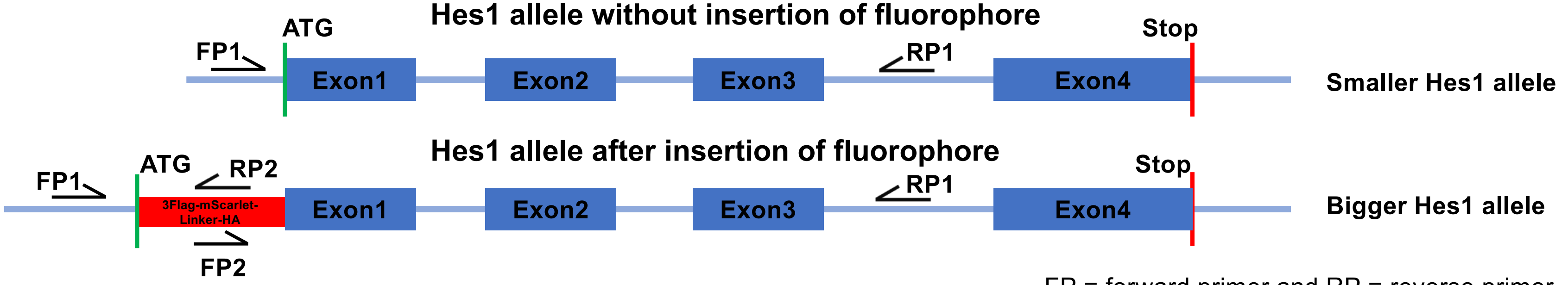


FP = forward primer and RP = reverse primer

(B)

**Genotyping summary of the additional CRISPR clones**

| Clone ID | Smaller Hes1 allele (without FF insertion) | Bigger Hes1 allele (with FF insertion) | Nature of the line |
|---|---|---|---|
| M18 | 17bp deletion near the PAM site, leading to premature STOP codon after aa 26 | mScarlet insertion at N terminal | Hemizygous |
| Cl12 | No mutations detected in the smaller Hes1 allele | mScarlet insertion at N terminal | Heterozygous |

FF = fluorophore, bp = base pair and aa = amino acid

Detailed sequencing data not shown. Primer details are available on request.

(C)

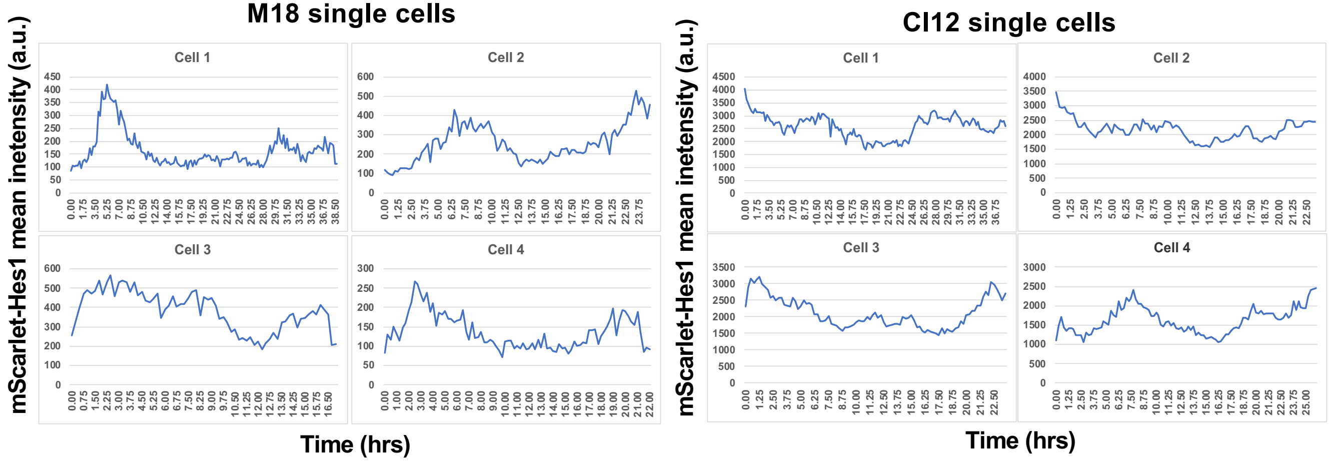


(D)

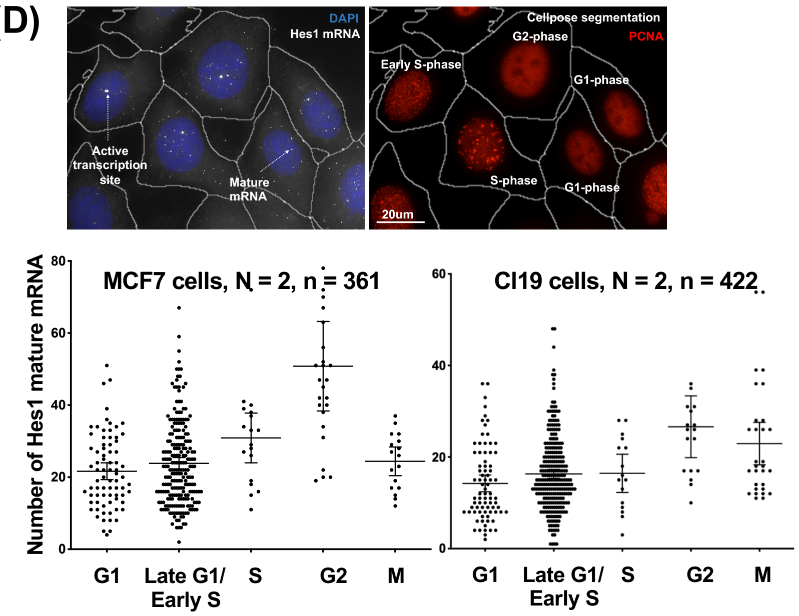


**Figure S4**

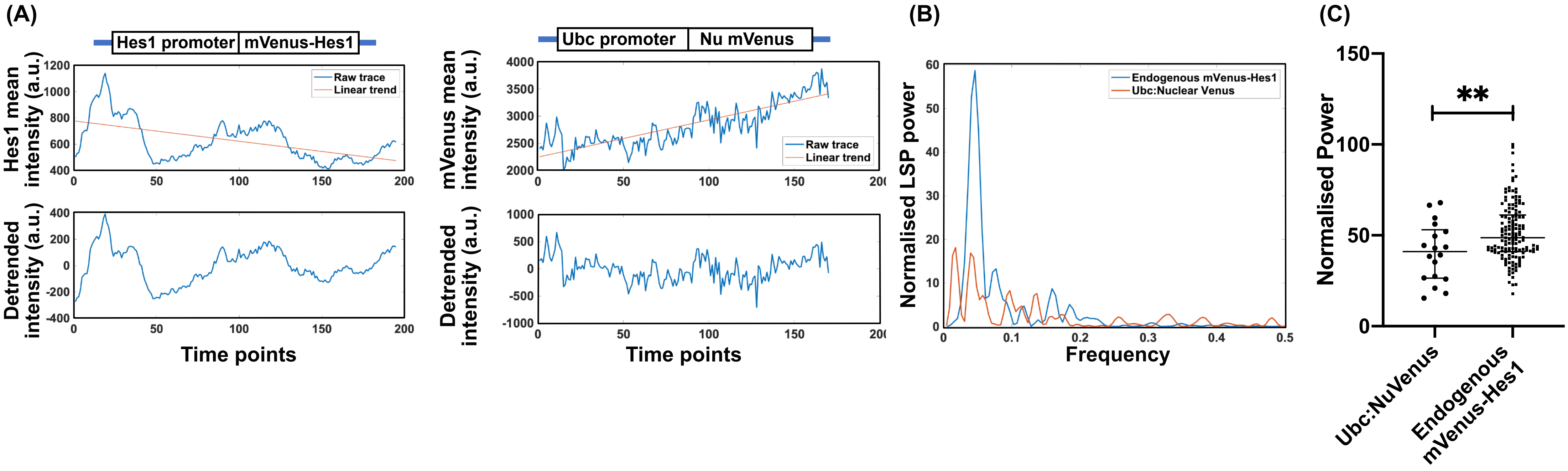
(A)
Hes1 promoter
mVenus-Hes1
Hes1 mean intensity (a.u.)
Raw trace
Linear trend
Detrended intensity (a.u.)
Time points
Ubc promoter
Nu mVenus
mVenus mean intensity (a.u.)
Raw trace
Linear trend
Detrended intensity (a.u.)
Time points
(B)
Normalised LSP power
Endogenous mVenus-Hes1
Ubc:Nuclear Venus
Frequency
(C)
Normalised Power
**
Ubc:NuVenus
Endogenous mVenus-Hes1

Figure S5

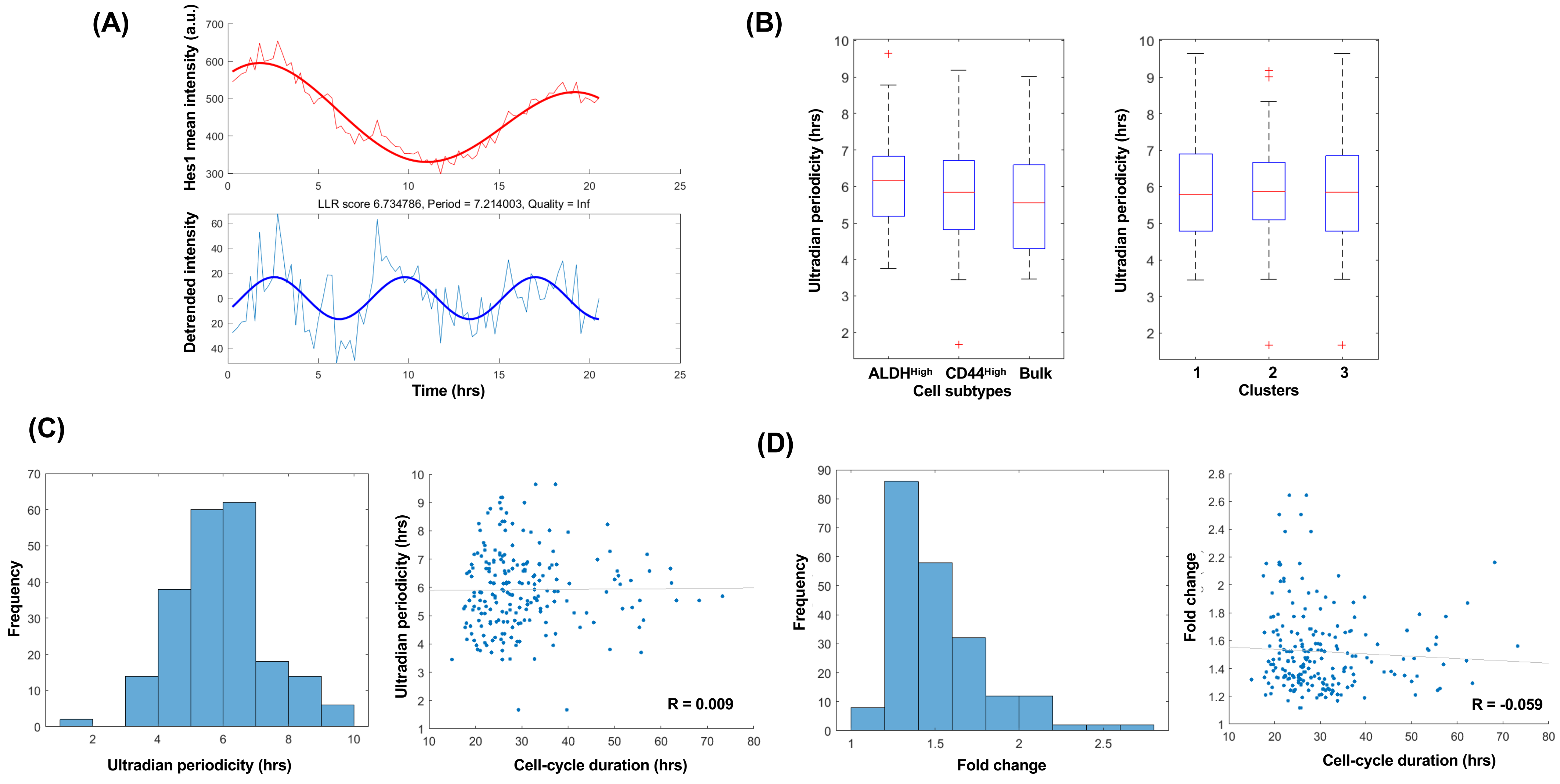
(A)
Hes1 mean intensity (a.u.)
LLR score 6.734786, Period = 7.214003, Quality = Inf
Detrended intensity
Time (hrs)
(B)
Ultradian periodicity (hrs)
ALDH^High CD44^High Bulk
Cell subtypes
Clusters
(C)
Frequency
Ultradian periodicity (hrs)
Cell-cycle duration (hrs)
R = 0.009
(D)
Frequency
Fold change
Cell-cycle duration (hrs)
R = -0.059

Figure S6

#### (A) Conversion of mVenus-Hes1 time-tracks into heat maps

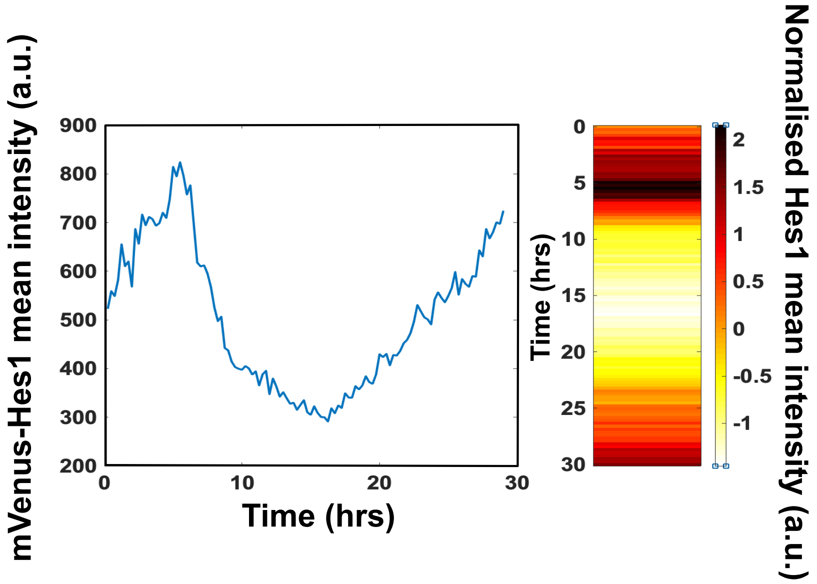


#### (B) Cell lineage tree

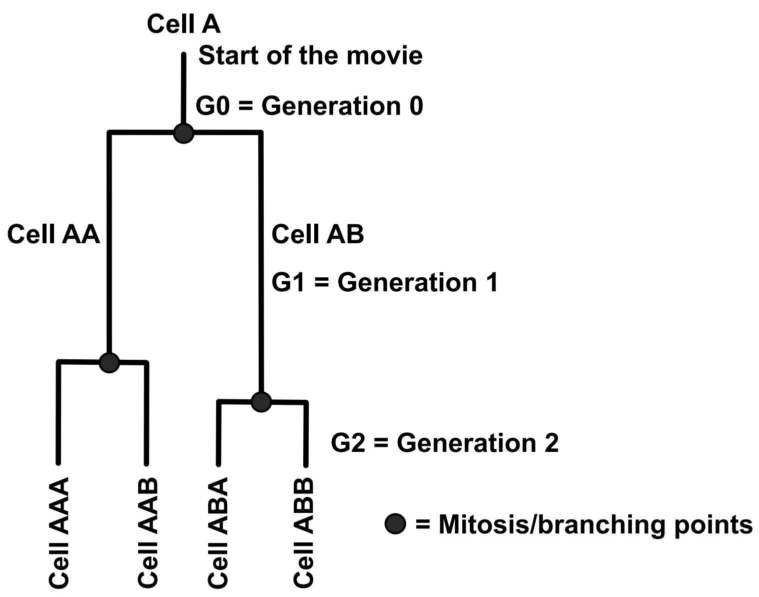


Figure S7

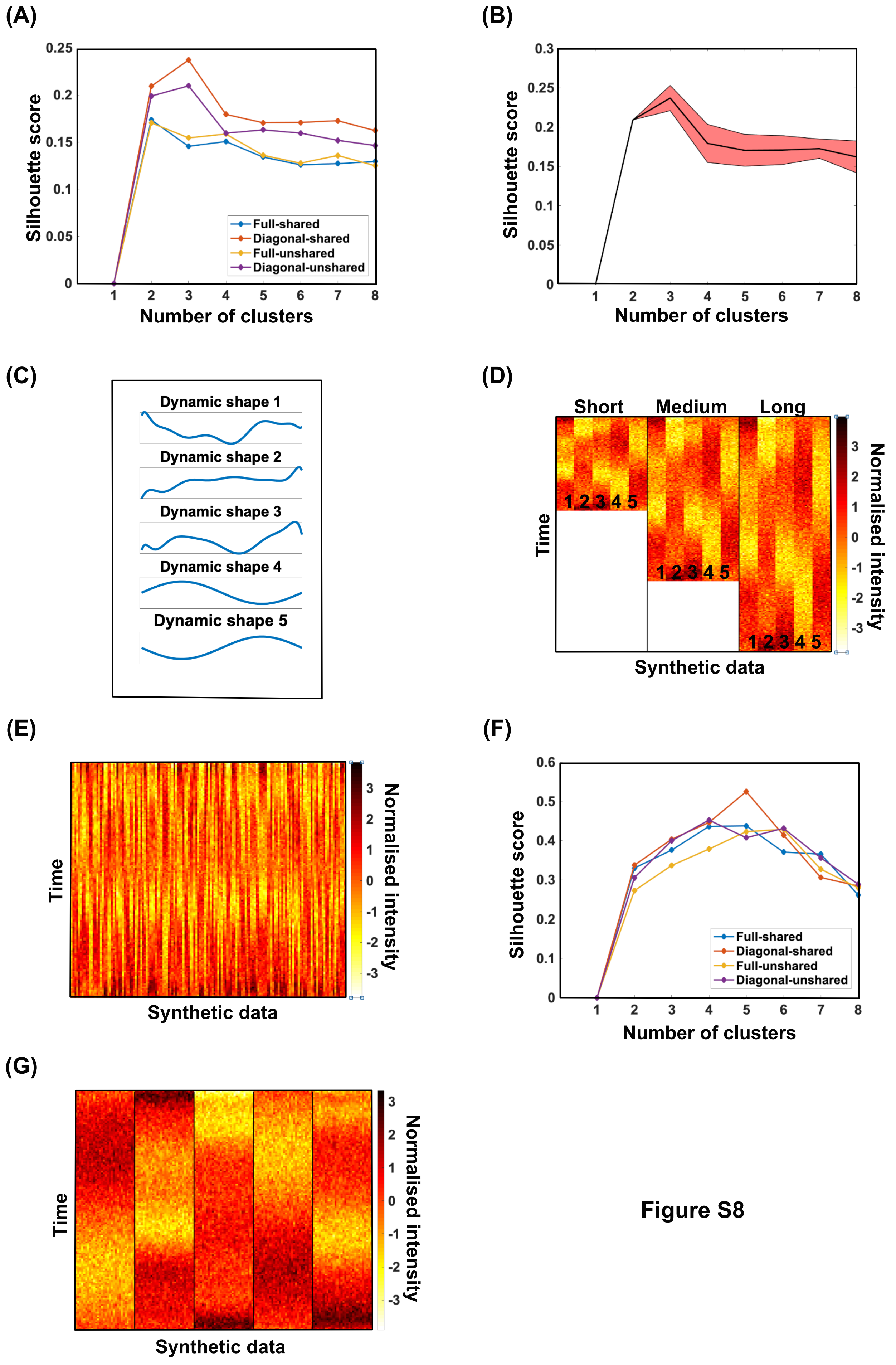
(A)
Silhouette score
Number of clusters
Full-shared
Diagonal-shared
Full-unshared
Diagonal-unshared
(B)
Silhouette score
Number of clusters
(C)
Dynamic shape 1
Dynamic shape 2
Dynamic shape 3
Dynamic shape 4
Dynamic shape 5
(D)
Short
Medium
Long
Time
1 2 3 4 5
Synthetic data
Normalised intensity
(E)
Time
Synthetic data
Normalised intensity
(F)
Silhouette score
Number of clusters
Full-shared
Diagonal-shared
Full-unshared
Diagonal-unshared
(G)
Time
Synthetic data
Normalised intensity

Figure S8

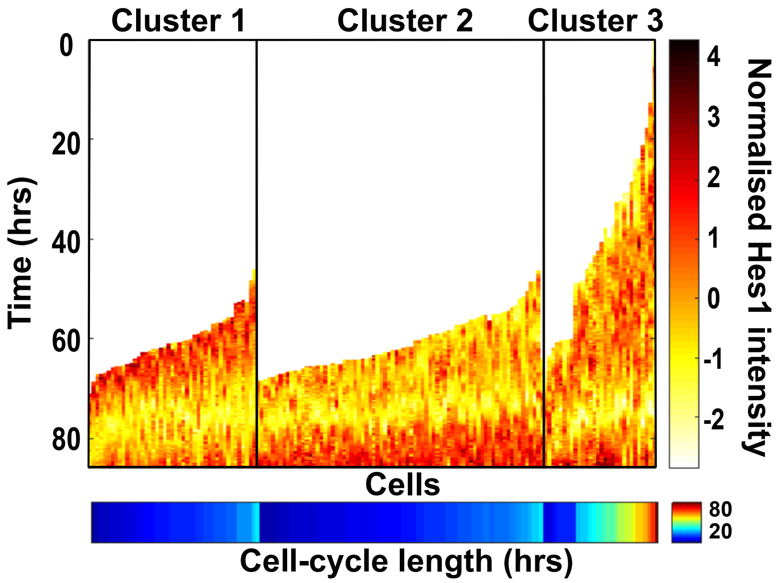
Cluster 1
Cluster 2
Cluster 3
0
20
40
60
80
Time (hrs)
4
3
2
1
0
-1
-2
Normalised Hes1 intensity
Cells
80
20
Cell-cycle length (hrs)

Figure S9

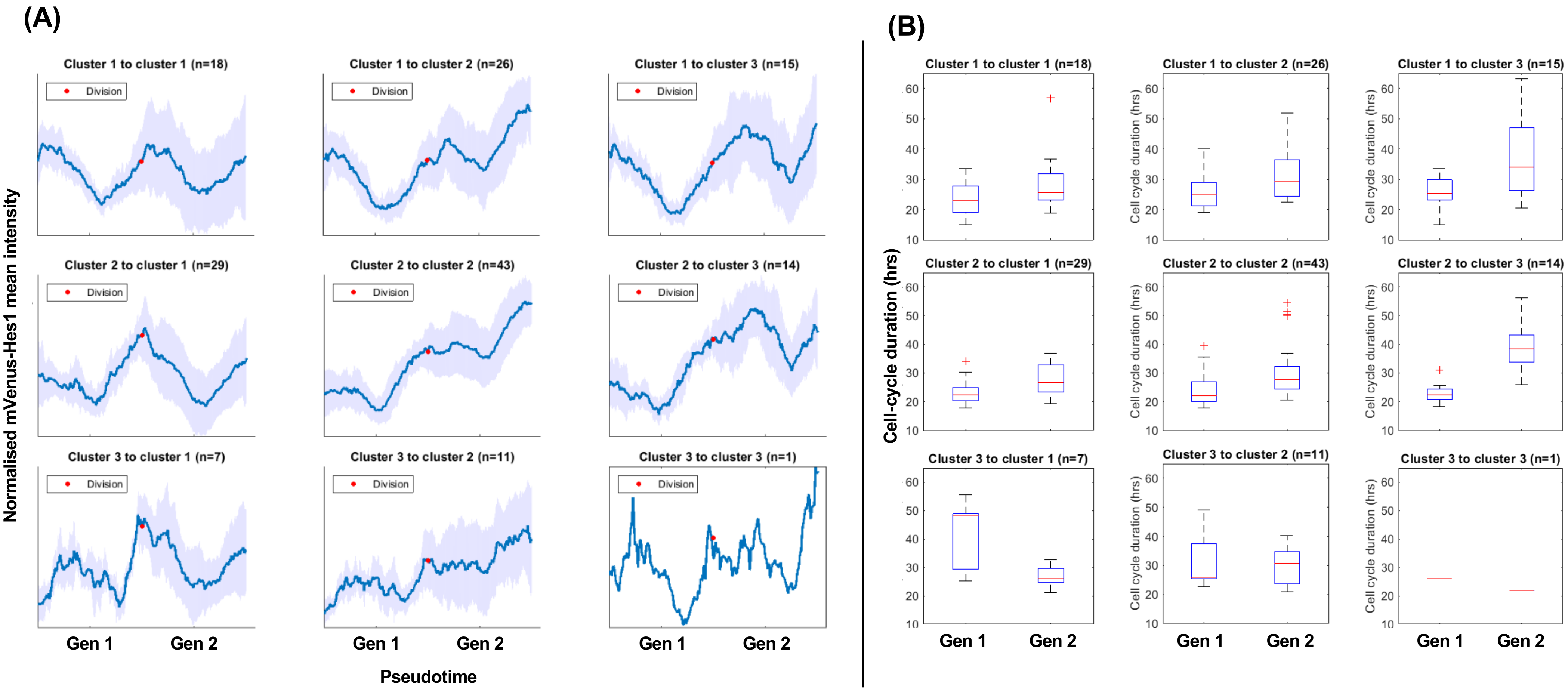

(A)
Normalised mVenus-Hes1 mean intensity
Cluster 1 to cluster 1 (n=18)
Cluster 1 to cluster 2 (n=26)
Cluster 1 to cluster 3 (n=15)
Cluster 2 to cluster 1 (n=29)
Cluster 2 to cluster 2 (n=43)
Cluster 2 to cluster 3 (n=14)
Cluster 3 to cluster 1 (n=7)
Cluster 3 to cluster 2 (n=11)
Cluster 3 to cluster 3 (n=1)
Division
Gen 1
Gen 2
Pseudotime
(B)
Cell-cycle duration (hrs)
Cell cycle duration (hrs)
Cluster 1 to cluster 1 (n=18)
Cluster 1 to cluster 2 (n=26)
Cluster 1 to cluster 3 (n=15)
Cluster 2 to cluster 1 (n=29)
Cluster 2 to cluster 2 (n=43)
Cluster 2 to cluster 3 (n=14)
Cluster 3 to cluster 1 (n=7)
Cluster 3 to cluster 2 (n=11)
Cluster 3 to cluster 3 (n=1)
10
20
30
40
50
60
Gen 1
Gen 2


**Figure S10**

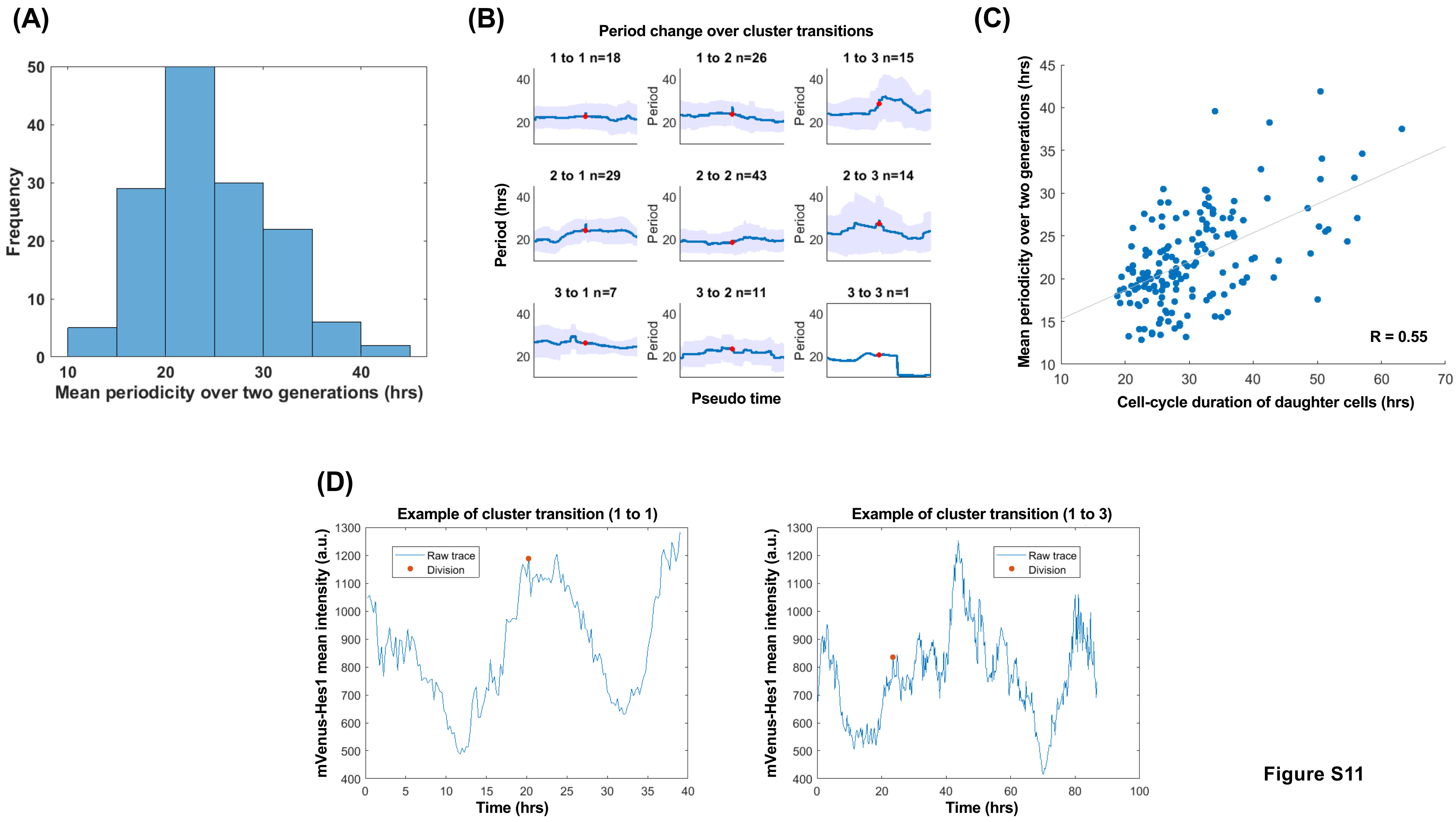
(A)
Frequency
Mean periodicity over two generations (hrs)
(B)
Period change over cluster transitions
1 to 1 n=18
1 to 2 n=26
1 to 3 n=15
2 to 1 n=29
2 to 2 n=43
2 to 3 n=14
3 to 1 n=7
3 to 2 n=11
3 to 3 n=1
Period (hrs)
Period
Pseudo time
(C)
Mean periodicity over two generations (hrs)
Cell-cycle duration of daughter cells (hrs)
R = 0.55
(D)
Example of cluster transition (1 to 1)
Example of cluster transition (1 to 3)
mVenus-Hes1 mean intensity (a.u.)
Raw trace
Division
Time (hrs)

Figure S11

(A)

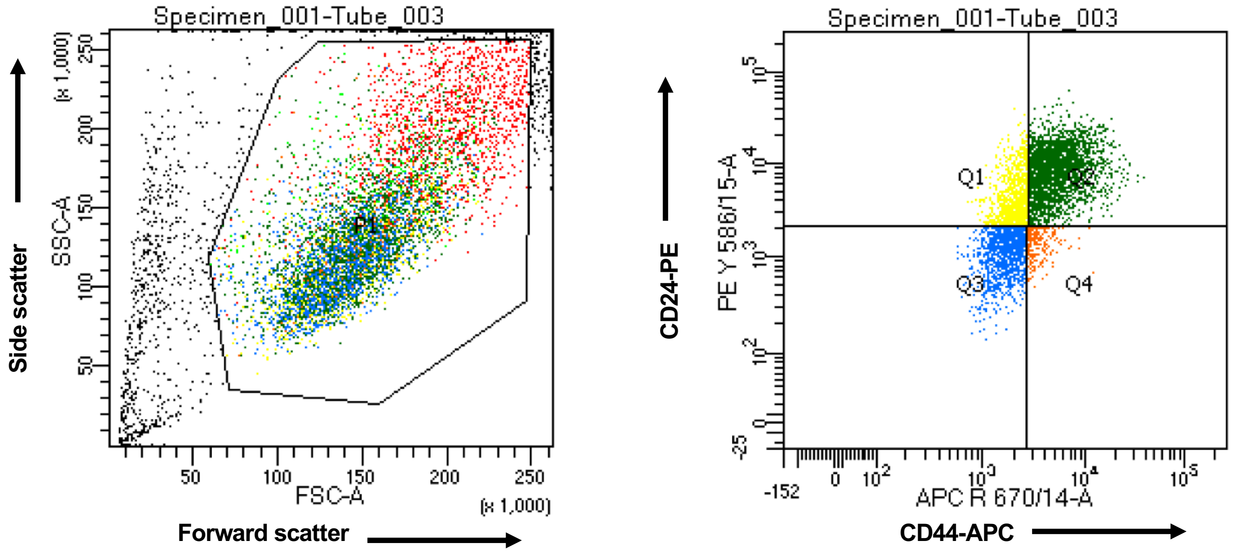


**FACS lineage tree and percentages of various cell populations**

Tube: Tube_003

| Population | #Events | %Parent | %Total |
|---|---|---|---|
| All Events | 10,000 | #### | 100.0 |
| P1 | 6,906 | 69.1 | 69.1 |
| P2 | 5,442 | 78.8 | 54.4 |
| P3 | 5,128 | 94.2 | 51.3 |
| Q1 | 994 | 19.4 | 9.9 |
| Q2 | 2,695 | 52.6 | 27.0 |
| Q3 | 1,193 | 23.3 | 11.9 |
| Q4 | 246 | 4.8 | 2.5 |

Q4 = $CD44^{High}CD24^{Low}$ population

(B)

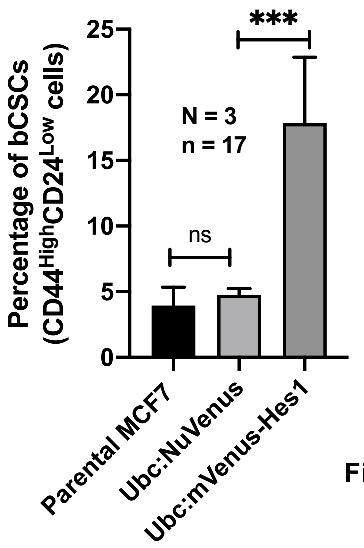


**Figure S12**

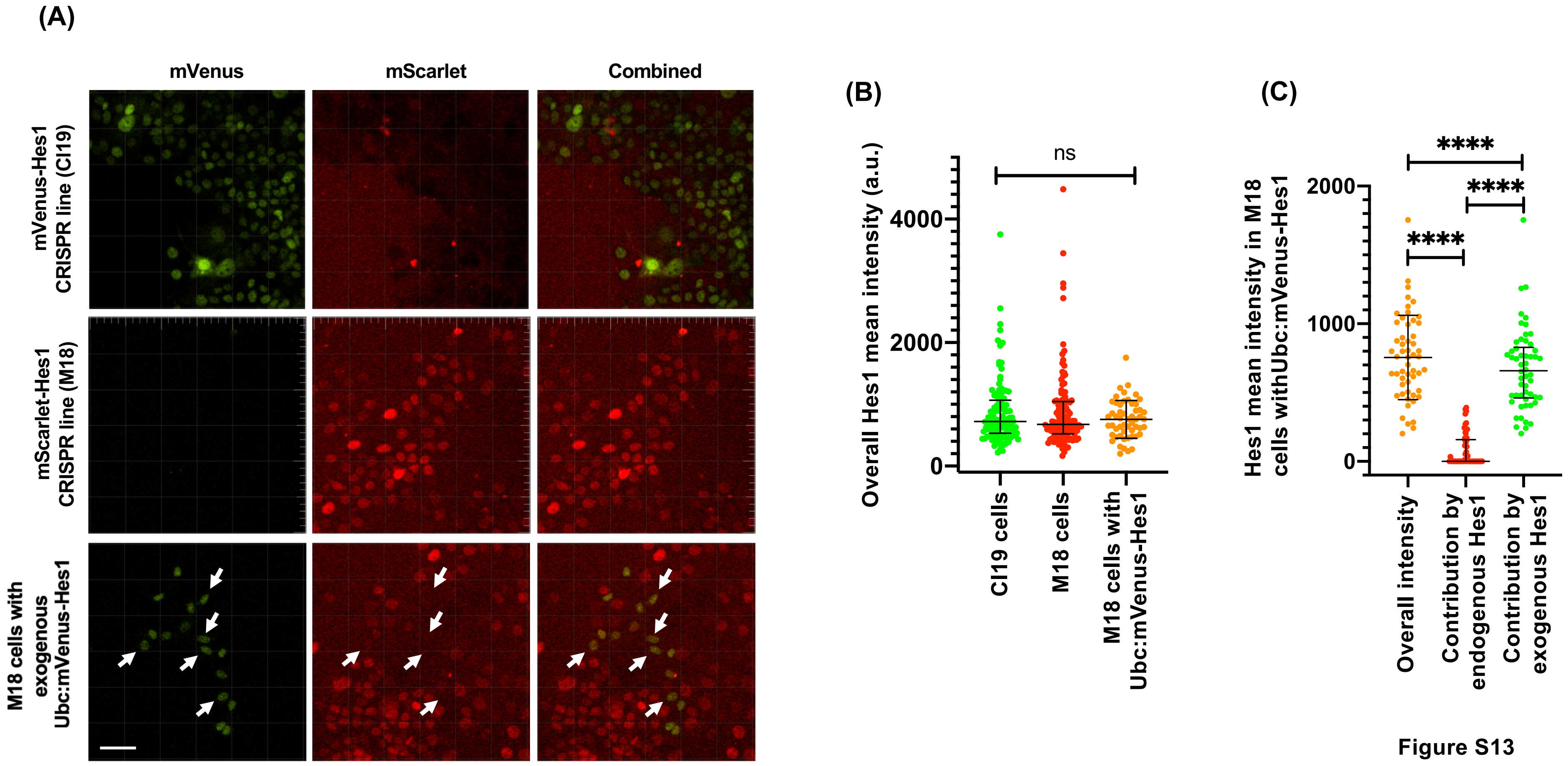

(A)
mVenus
mScarlet
Combined
mVenus-Hes1 CRISPR line (Cl19)
mScarlet-Hes1 CRISPR line (M18)
M18 cells with exogenous Ubc:mVenus-Hes1
(B)
Overall Hes1 mean intensity (a.u.)
ns
4000
2000
0
Cl19 cells
M18 cells
M18 cells with Ubc:mVenus-Hes1
(C)
Hes1 mean intensity in M18 cells withUbc:mVenus-Hes1
****
****
****
2000
1000
0
Overall intensity
Contribution by endogenous Hes1
Contribution by exogenous Hes1


Figure S13

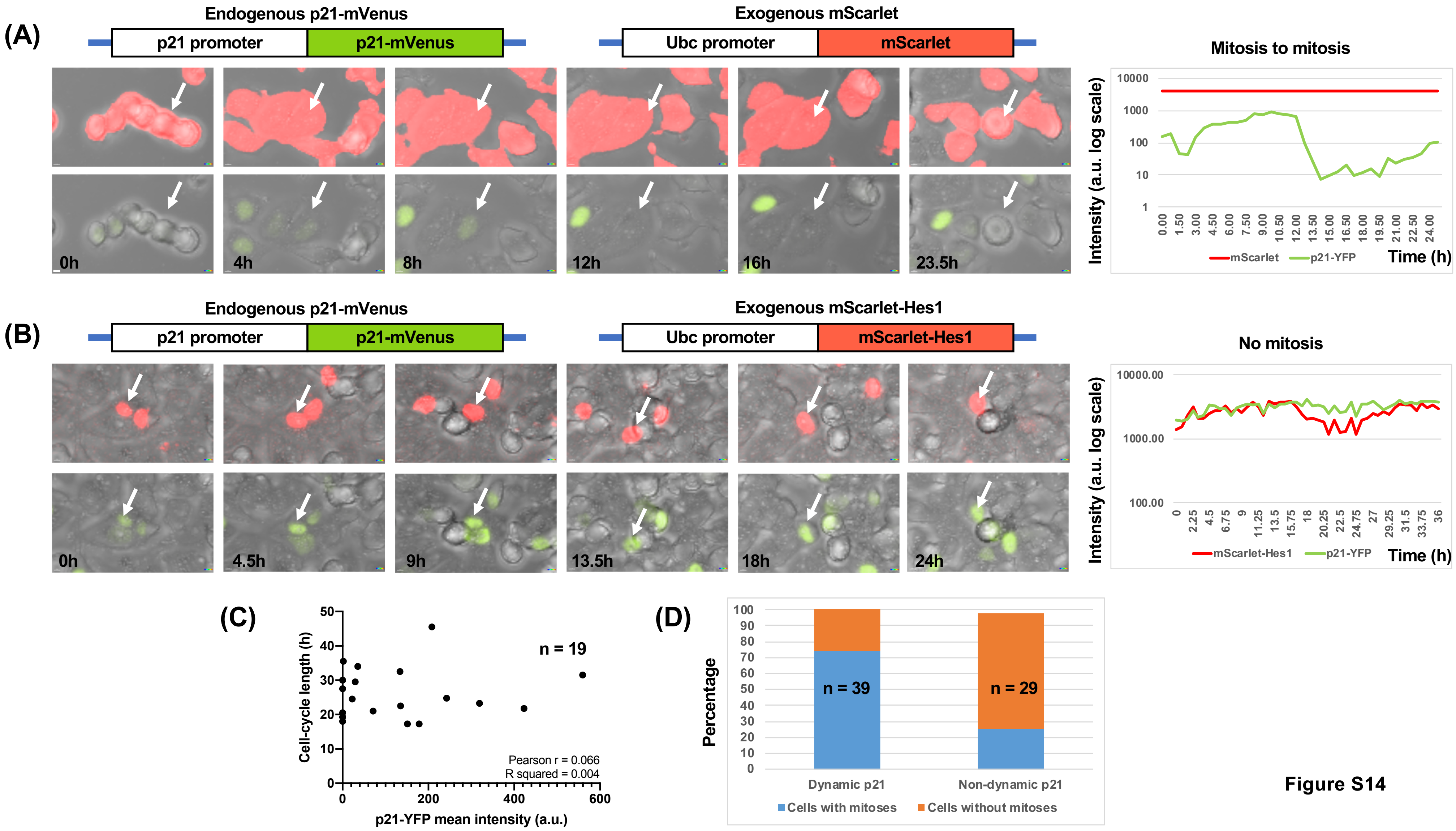

(A)
Endogenous p21-mVenus
p21 promoter
p21-mVenus
Exogenous mScarlet
Ubc promoter
mScarlet
0h
4h
8h
12h
16h
23.5h
Mitosis to mitosis
Intensity (a.u. log scale)
10000
1000
100
10
1
0.00
1.50
3.00
4.50
6.00
7.50
9.00
10.50
12.00
13.50
15.00
16.50
18.00
19.50
21.00
22.50
24.00
mScarlet
p21-YFP
Time (h)
(B)
Endogenous p21-mVenus
p21 promoter
p21-mVenus
Exogenous mScarlet-Hes1
Ubc promoter
mScarlet-Hes1
0h
4.5h
9h
13.5h
18h
24h
No mitosis
Intensity (a.u. log scale)
10000.00
1000.00
100.00
0
2.25
4.5
6.75
9
11.25
13.5
15.75
18
20.25
22.5
24.75
27
29.25
31.5
33.75
36
mScarlet-Hes1
p21-YFP
Time (h)
(C)
Cell-cycle length (h)
50
40
30
20
10
0
n = 19
Pearson r = 0.066
R squared = 0.004
0
200
400
600
p21-YFP mean intensity (a.u.)
(D)
Percentage
100
90
80
70
60
50
40
30
20
10
0
n = 39
n = 29
Dynamic p21
Non-dynamic p21
Cells with mitoses
Cells without mitoses


Figure S14
